# Schizophrenia-associated variation at *ZNF804A* correlates with altered experience-dependent dynamics of sleep slow-waves and spindles in healthy young adults

**DOI:** 10.1101/2020.05.01.072165

**Authors:** Ullrich Bartsch, Laura J Corbin, Charlotte Hellmich, Michelle Taylor, Kayleigh E Easey, Claire Durant, Hugh M Marston, Nicholas J Timpson, Matthew W Jones

**Affiliations:** School of Physiology, Pharmacology & Neuroscience, University of Bristol, Bristol, UK; Translational Neuroscience, Eli Lilly & Co Ltd UK, Erl Wood Manor, Windlesham, UK; MRC Integrative Epidemiology Unit, University of Bristol, Bristol, UK; Clinical Research and Imaging Centre (CRIC), University of Bristol, Bristol, UK

**Keywords:** psychosis, schizophrenia, genetics, sleep, slow wave, spindle, motor sequence task, ALSPAC

## Abstract

**Background:** The rs1344706 polymorphism in *ZNF804A* is robustly associated with schizophrenia (SZ), yet brain and behavioral phenotypes related to this variant have not been extensively characterized. In turn, SZ is associated with abnormal non-rapid eye movement (NREM) sleep neurophysiology. To examine whether rs1344706 is associated with intermediate neurophysiological traits in the absence of disease, we assessed the relationship between genotype, sleep neurophysiology, and sleep-dependent memory consolidation in healthy participants.

**Methods:** We recruited healthy adult males, with no history of psychiatric disorder, from the Avon Longitudinal Study of Parents and Children (ALSPAC) birth cohort. Participants were homozygous for either the SZ-associated ‘A’ allele (N=25) or the alternative ‘C’ allele (N=22) at rs1344706. Actigraphy, polysomnography (PSG) and a motor sequencing task (MST) were used to characterize daily activity patterns, sleep neurophysiology and sleep-dependent memory consolidation.

**Results:** Average MST learning and sleep-dependent performance improvements were similar across genotype groups, but with increased variability in the AA group. CC participants showed increased slow-wave and spindle amplitudes, plus augmented coupling of slow-wave activity across recording electrodes after learning. Slow-waves and spindles in those with the AA genotype were insensitive to learning, whilst slow-wave coherence decreased following MST training.

**Conclusion:** We describe evidence that rs1344706 polymorphism in *ZNF804A* is associated with changes in experience- and sleep-dependent, local and distributed neural network activity that supports offline information processing during sleep in a healthy population. These findings highlight the utility of sleep neurophysiology in mapping the impacts of SZ-associated variants on neural circuit oscillations and function.

## INTRODUCTION

Schizophrenia (SZ) is a debilitating psychiatric disorder with a lifetime prevalence of up to 4% (1). SZ etiology is complex and heterogenous, but an estimated heritability of up to 80% reflects critical genetic contributions to SZ liability (2,3). Recent efforts in cataloguing the genetic architecture of SZ have generated a list of over 100 loci thought to contribute in some way to the development of the disease (4,5). Despite most of these risk variants having small individual effects and acting in combination with other genetic and environmental factors, elucidating the neuronal changes downstream of genetic liability remains crucial for understanding normal brain development and the etiology of psychiatric disorders.

The single nucleotide polymorphism (SNP) rs1344706 within the second intron of *ZNF804A* was the first SNP to show genome-wide significant association for psychosis diagnosed in both bipolar disorder and SZ (6). This finding has been replicated in subsequent genome wide association studies (GWAS) (4,7–9) including a fine-mapping study which confirmed rs1344706 as the most strongly associated variant at the locus, with an OR for SZ of 1.10 [1.07 – 1.14] (10). *ZNF804A* is expressed in the brain and is predicted to encode a protein with a C2H2 zinc finger domain, indicating a role in transcriptional regulation (8,10) and thus likely complex biological functions (11). rs1344706 has been linked to a number of behavioral and neuronal phenotypes (12,13), correlating with altered neuroanatomy (14,15) (but see (16) for a null result), abnormal neurophysiology (17–19) and cognitive phenotypes (20–22). In particular, *ZNF804A* genotype has been associated with cortico-hippocampal functional connectivity in healthy control subjects (23,24) and also in SZ patients and their unaffected siblings (17,25).

Whilst cognitive deficits are an established feature of SZ (26,27), links have recently been made between cognitive symptom dimensions and abnormal sleep. Sleep disturbances are a core feature of SZ (28,29) and include increased sleep latency and decreased total sleep time, independent of neuroleptic treatment (30). At the level of neural network activity, sleep in patients also features changes in characteristic electroencephalography (EEG) oscillations, particularly during NREM sleep. Thalamo-cortical spindle oscillations are a defining feature of NREM and are reduced in patients with SZ (31–34). Consistent with the roles of spindle oscillations in memory consolidation in healthy participants (35–38) spindle deficits in SZ have been linked to cognitive deficits in patients (39,40). More recently, slow oscillations and their coordination with spindles have also been implicated in contributing to deficits in sleepdependent memory consolidation in patients (41–43).

Overall, there is convergent evidence that circuit abnormalities in SZ are reflected by changes in sleep physiology that, in turn, may be important for cognitive symptoms (44). In principle, linking specific genetic variations with sleep neurophysiology phenotypes holds the promise of illuminating a broader understanding of genetic effects and potential mechanisms of neural circuit dysfunction in SZ. Here we used a recall-by-genotype approach (45) to recruit healthy individuals who were homozygous at rs1344706 in order to reduce the issues of confounding and reverse causality that commonly effect traditional observational case/control studies. In this case, the availability of genetic data in a large and engaged cohort study allowed for efficient and balanced recruitment of participants into a detailed examination of sleep architecture and neurophysiology. We aimed to test the hypothesis that, in the absence of disease, rs1344706 genotype would associate with the facets of abnormal sleep neurophysiology and sleep-dependent memory consolidation seen in SZ.

## METHODS AND MATERIALS

The study design and protocol was published in advance in (46); raw and processed data and metadata are available upon application to the Avon Longitudinal Study of Parents and Children (ALSPAC) Executive Committee through a standard application process (see http://www.bristol.ac.uk/alspac/researchers/access/).

### Participants

Healthy males aged 21-23 years and of European ancestry were recruited from ALSPAC, a prospective birth cohort allowing the study of health and development across the life course (47,48). Participants were invited based on homozygosity either for the rs1344706 allele previously associated with increased liability for SZ (AA group), or for the alternative allele (CC group).

Ethical approval for the study was obtained from the ALSPAC Ethics and Law Committee (ref. 9224). The data collection protocol was previously approved by The University of Bristol Faculty of Science Human Research Ethics Committee as part of a pilot study (ref. 8089). All participants provided informed consent to participate in the study following the recommendations of the ALSPAC Ethics and Law Committee. For a detailed description of the cohort and recruitment see Supplemental Methods.

### Procedures

Data were collected from each participant over approximately two weeks, beginning and ending with a night of polysomnography (PSG, including 9-channel EEG) at the Clinical Research and Imaging Centre at the University of Bristol (Figure 1). Both researchers and participants were blind to participant genotype throughout data collection. During visits, participants completed sleep-based questionnaires in order to assess self-rated sleep quality and collect information about subjective experience of their night in the sleep laboratory (see a detailed description of all procedures in Supplemental Procedures).

**Figure 1:**
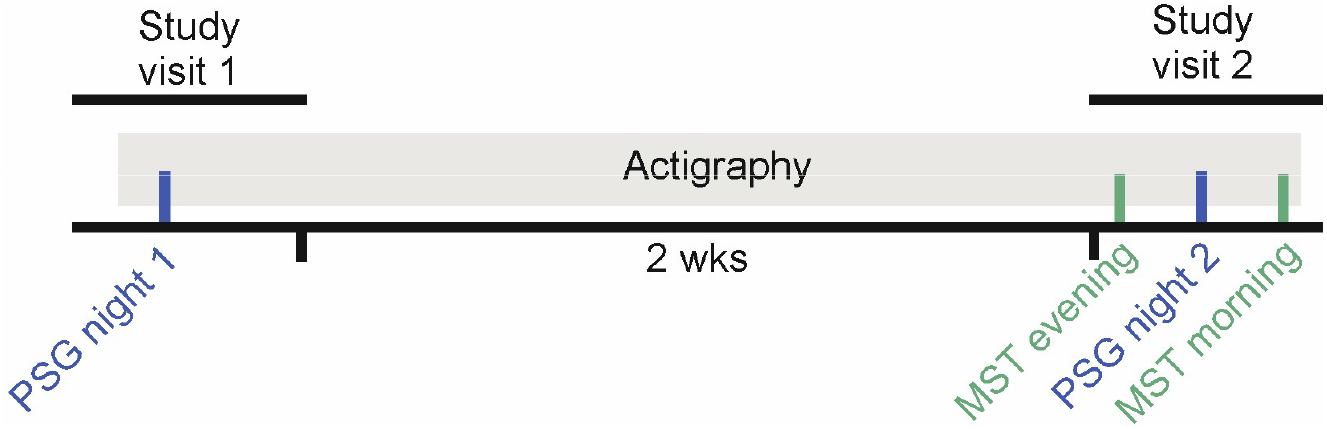
Study design. The study included two study visits to the sleep lab at the Clinical Research & Imaging Centre in Bristol with two weeks of actigraphy monitoring between visits. Participants visited the sleep lab for a baseline polysomnography (PSG) recording night on their first visit (night 1). They were also issued an actigraphy watch which they wore until the end of the second study visit. During the second visit, participants were trained on the motor sequence task (MST) in the evening and tested in the morning, with an intervening second PSG recording (night 2).

### Analysis

#### Behavioral data

All paper questionnaires were manually scored and transcribed to spreadsheets. MST performance was quantified using the number of correct sequences and the reaction time during correct sequences. These variables were used to calculate average values for trial 10-12 of the evening session (‘training performance’) and average values for trials 1-3 of the morning session (‘test performance’). Overnight improvement was calculated as the percentage change in each outcome measure from training to test (49). Actigraphy data were manually annotated in MotionWare (CamNtech, UK) to derive sleep architecture and circadian rhythm measures.

#### Polysomnography data

Polysomnography was scored by an experienced expert (blinded to participant genotype) based on AASM criteria (50) using REMLogic software (Natus Europe GmbH, Germany). Sleep architecture was quantified using standard variables including total sleep time (TST) and sleep onset latency (SOL) (see Polysomnography Analyses in Supplemental Methods). EEG traces were analyzed using automatic detection of characteristic NREM sleep events - slow waves (SW), delta waves, slow and fast spindle events as described earlier (43,51). Characteristic NREM events were further characterized using multitaper spectra and coherence using the Chronux toolbox (www.chronux.org).

### Statistical methods

A detailed description of all analysis and statistical methods can be found in Supplemental Methods; Tables S1, S2 show a full record of methods and their alignment to analytical arguments. In brief, behavioral measures were analyzed either by a comparison of means across groups (two-sample two-sided t-test or Wilcoxon rank-sum test) or by fitting a linear mixed model with genotype and MST session (training versus testing) fitted as fixed effects (using Stata v14.2 (52)). PSG-derived sleep architecture measures were analyzed in a linear mixed model with genotype and recording night (night 1: baseline, night 2: learning) fitted as fixed effects. PSG-derived event properties, power and coherence measures were compared across genotype groups, electrodes, recording nights (night 1: baseline, night 2: learning) and sleep stages (N2, N3) using a linear mixed model framework and a stepwise reduction procedure implemented using the lme4 (53) and lmerTest (54) packages in R. We built a full model of the general form [y ~ *genotype + night + electrode + sleep_stage + (genotype * night) + (1 (ID)*], where y is any derived sleep variable, and then applied backward elimination of non-significant model terms using the R function step which is part of the R package lmerTest (54). Results presented are mean ± standard error (SE) unless stated otherwise.

## RESULTS

Data were collected from 47 participants (25 AA and 22 CC). The two genotype groups did not differ in maternal education, social class, psychosis-like symptoms at age 18, or in the Wechsler Abbreviated Scale of Intelligence at age 15 (Table S3). Data from seven participants were excluded due to missing or corrupted data (see Supplemental Results for further explanation); we therefore present results for 40 participants (Figure S1).

### Increased variability in Motor Sequence Task performance in the rs1344706 AA group

Overall performance levels for practice-dependent increases in the number of correct sequences (NCS) – and corresponding decreases in button press latency within correct sequences (‘reaction time’, RT) – were comparable between genotype groups. Figure 2, panels A, D show the MST learning curves for both genotype groups and Figure 2, panels B, E show the averages of the last 3 trials in the evening and first 3 trials in the morning that are used to calculate overnight improvement. Participants in both groups improved overnight in mean NCS (overnight change in NCS, CC: 16.9% with standard deviation (SD): 9.6, AA: 15.9% SD: 16.8, Figure 2C, Table 1) and RT (overnight change in absolute RT, CC: 10.5% (SD: 6.2), AA: 8.3% (SD 11.9), Figure 2F, Table 1).

**Figure 2:**
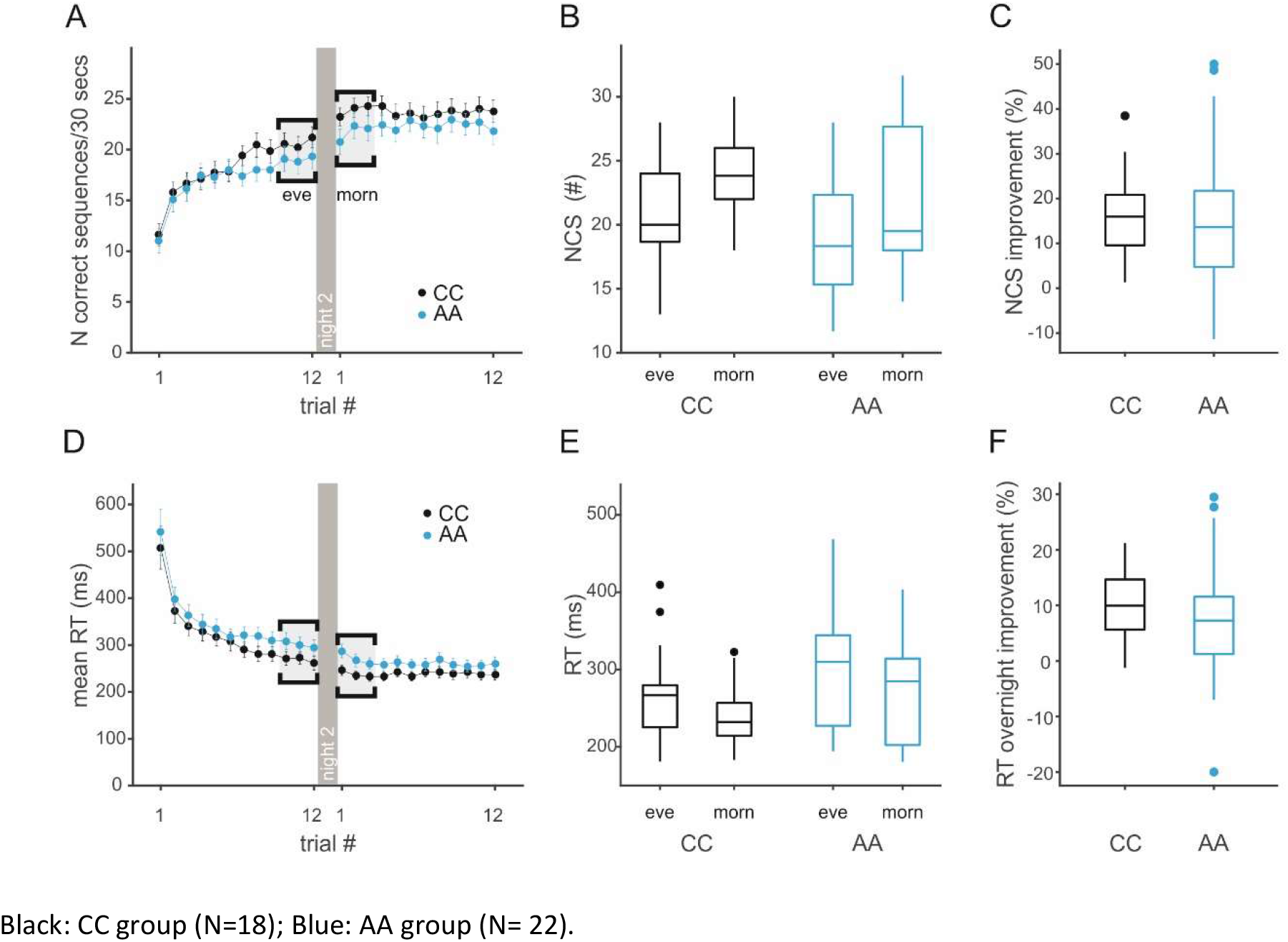
Average MST learning curves and overnight improvement by genotype group. A) MST learning curves showing the number of correct sequences per trial. Night 2 is indicated by a dark grey separator, the last 3 and first 3 trials used to calculate the average for the evening (eve) and morning (morn) performance are highlighted in light grey. B) Box plot showing median number of correct sequences (last 3 trials in the evening v first 3 trials in the morning) for each MST session and genotype group. (Plots indicate the median, with boxes showing the 25th and 75^th^ percentile of data, whiskers indicate the range of values inside 1.5* interquartile range, extreme values (outside 1.5 IQR) are plotted as individual data points, see Statistical Methods for details). C) Boxplot showing the median of overnight improvement in number of correct sequences/30 s trial as percentage change from evening to morning performance. D) Learning curves as in A but for the mean reaction time (RT, button press latency within a correct sequence) per trial. E) Boxplot showing the median RT during correct sequence button presses (last 3 trials in the evening v first 3 trials in the morning) for each MST session and genotype group. F) Boxplot of median overnight improvement in RT measured as absolute percentage change from evening to morning performance

Linear mixed modelling of the MST performance data confirmed effects of session (training vs. test) on NCS (session: F (1, 39)= 79.1, p = 6.38e-11) and RT (session: F (1, 39)= 28.8, p = 3.93e-06, Table 2), suggesting sleep-dependent consolidation of motor memory in both genotype groups. There was no strong evidence for an effect of genotype on task performance, but point estimates suggested that AA group participants produced fewer correct sequences (F(1, 38) = 1.61, *p* = 0.21) and had slower reaction times (F (1, 38)= 3.0, *p* = 0.09, Table 2).

Interestingly, the AA group showed higher variance in overnight improvement in NCS (SD CC: 9.6, AA: 16.8, two-sample variance comparison *p* = 0.02, Table 1) and RT (SD CC: 6.2, AA: 11.9, two-sample variance comparison *p* = 0.01). This higher variance was particularly pronounced during the morning test session (SD NCS, CC: 3.6, AA: 5.5, Levene’s test *p* = 0.02; SD RT, CC: 41 ms, AA: 62 ms, Levene’s test *p* = 0.02).

### Sleep timing, architecture and quality appear unaffected by rs1344706 genotype

We found no evidence for consistent effects of genotype on diurnal rhythmicity (Table S4), sleep architecture derived from the PSG (Figure 3 and Table S5) or subjective/objective sleep quality (Table S6).

**Figure 3:**
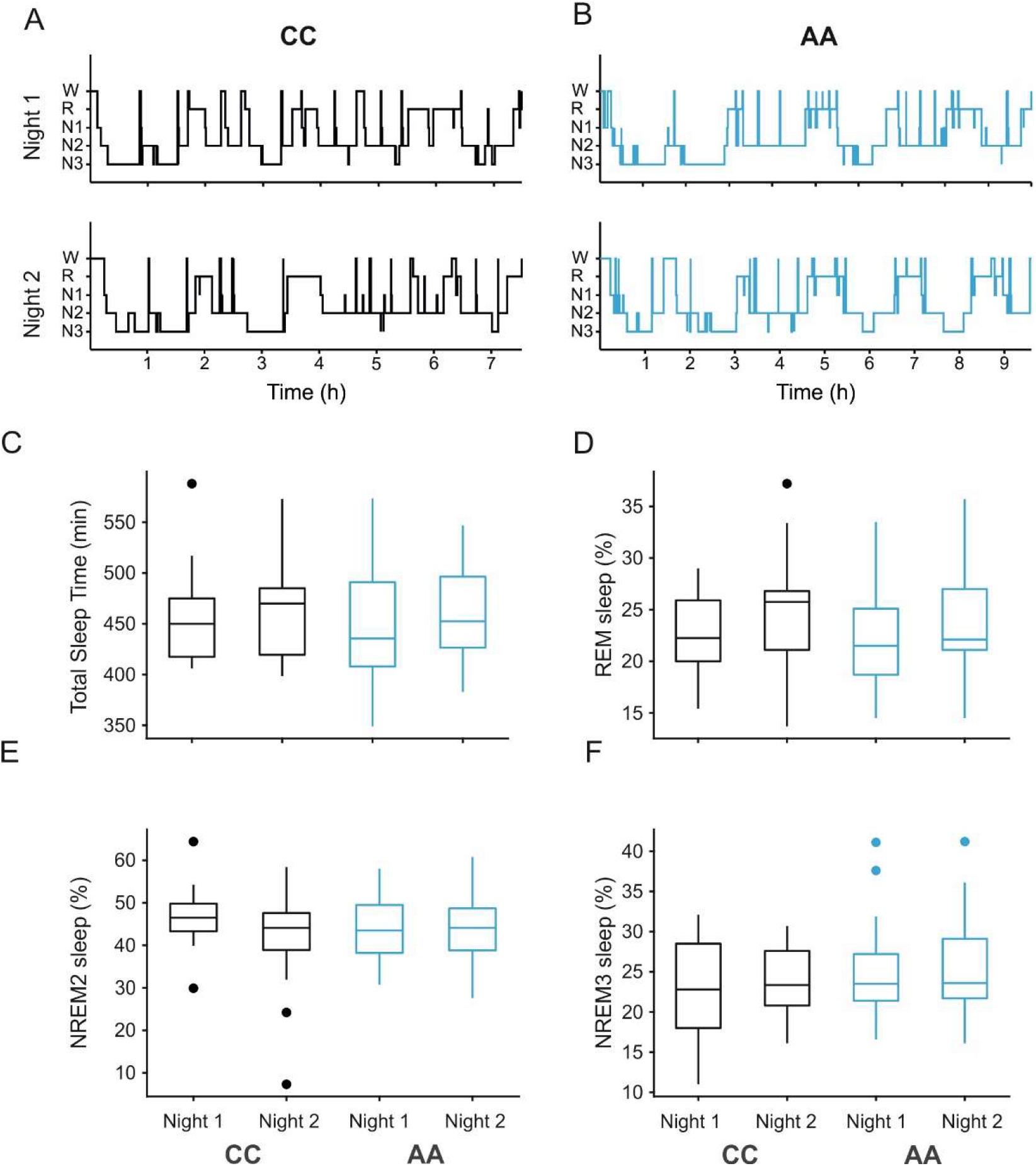
Example hypnograms and sleep architecture variables from polysomnography. A) Hypnogram examples from a CC individual for nights 1 and 2, x- axis shows time in hours from lights off. Sleep stages are Wake (W), REM (R), NREM1 (N1, NREM2 (N2), NREM3 (N3). B) Hypnogram examples from an AA individual for nights 1 and 2, x- axis shows time in hours from lights off C-F) Boxplots of PSG derived variables of sleep architecture by genotype group (CC, black; AA, blue) and recording night. Outlier (outside the 25^th^ and 75^th^ percentile) are included as individual data points (o symbol). We observed no significant effects of night or genotype using linear mixed model approach for C) Total Sleep Time (min), D) Percentage of REM sleep, E) Percentage of stage 2 NREM sleep or F) Percentage of stage 3 NREM sleep.

### rs1344706 genotype is associated with experience dependent changes in NREM sleep EEG oscillations

To assess potential neurophysiological correlates of variance in MST performance, we used custom detection algorithms to extract slow and delta waves (0.5-4Hz), and slow (9-12Hz) and fast spindles (13-16Hz). Figure 4, panels A and B show wave-triggered averages, revealing the morphologies of slow waves and fast spindles recorded at electrodes F3 and Cz: waveforms appeared similar across genotype group and recording night at these electrode locations. Panels C and D show average extracted amplitude values for each event at the electrode locations F3 and Cz. (For a complementary spectral analysis of power in corresponding frequency bands see Supplementary Results and Tables S7, S8.)

**Figure 4:**
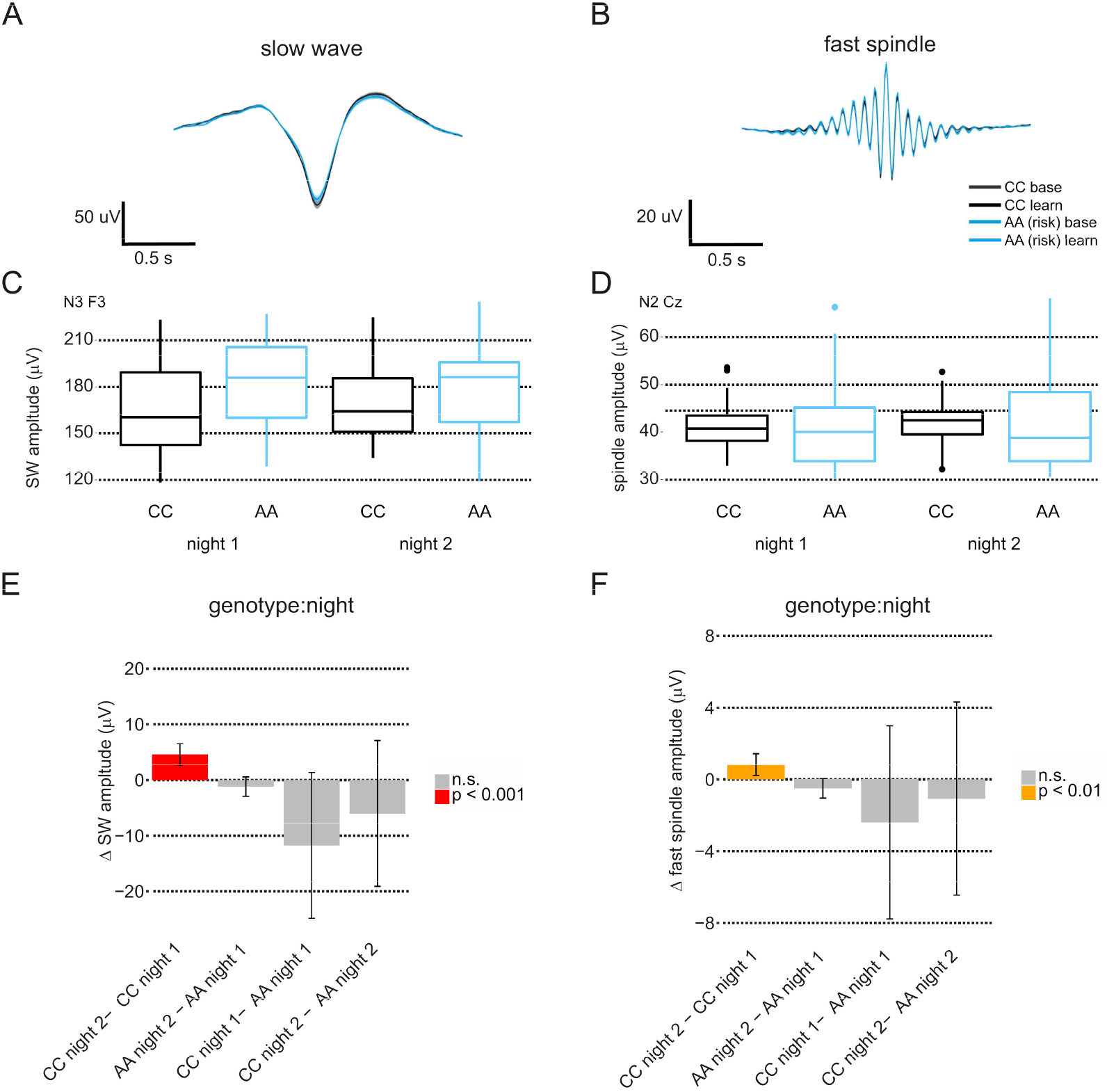
Slow wave and spindle properties. A) SW wave triggered average at F3 for all genotype groups and both recording nights. CC, baseline night, grey, CC, learning night, black, AA baseline night, light blue, AA learning night, dark blue. We found no significant difference for any time bin (p<0.05, Wilcoxon ranksum test, no correction). B) Spindle wave triggered average at Cz for all genotype groups and both recording nights, same colors as in A. We found no significant difference for any time bin (p<0.05, Wilcoxon ranksum test, no correction). C) Estimated marginal means for SW amplitude values for SW recorded at F3, during N3 sleep by genotype group and by recording night. D) Estimated marginal means of fast spindle amplitude values for spindles recorded at Cz, during N2 sleep for both genotypes and both recording nights. E) Estimated marginal means differences for the factors genotype and night estimated from a linear mixed model analysis of all detected SW amplitudes (using the R package lmerTest (see **Supplemental Methods** for details)). CC individuals show an increase in SW amplitude in night 2, but AA’s do not. Error bars indicate 95% confidence intervals. F) Same as E for all detected fast spindle amplitudes. Like SW amplitudes, CC individuals show an increase in fast spindle amplitude in night 2, but AA’s do not. Error bars indicate 95% confidence intervals.

#### Slow-wave amplitudes depend on experience, but only in the CC group

We did not detect differences in density of slow-wave events between nights or genotypes (Table S9). However, a linear mixed model analysis of slow-wave amplitudes from N2 and N3 sleep from all electrode locations suggested a main effect of night and an important interaction term (night by genotype) in the initial full model. After stepwise reduction, the night by genotype interaction remained (F(1, 1356.05) = 18.67, *p*= 1.67e-05, Table S9). Figure 4E shows the differences in estimated marginal means between nights, demonstrating an increase in slow-wave amplitudes from night 1 (baseline) to night 2 (learning) in CC participants (night 1: 100±4.80μV; night 2: 105.12±4.78μV, *p* <0.001, Table S10). In contrast, slow-wave amplitudes in the AA group remained similar across nights (night 1: 112±4.34μV, night 2: 111±4.34μV, n.s.). These SW event results were supported by very similar results for delta wave event properties (Tables S11, S12) and separate Fourier analyses of 0.5-1.5Hz power, which showed the same pattern of experience and genotype-dependent changes (Tables S7 and S8). Collectively, these analyses suggest that the coordinated firing of cortical populations during SW events may be modulated following learning in a genotype-dependent manner.

#### Spindle properties depend on experience, with differential effects of genotype

A linear mixed model analysis of 9-12Hz (slow) spindle event properties revealed night and genotype dependent associations with amplitude (Table S13), with a trend for an increase after learning in the CC group (31.1±2.0 μV during night 1 vs. 31.7±2.0 μV during night 2, *p*=0.05), but a decrease in the AA group (from 33.8±1.8 μV to 33.2±1.8 μV, *p*=0.03, Table S14). We also observed a main effect for night-dependent associations with slow spindle frequency, with small decreases in slow spindle frequency after motor learning in both groups (from 11.36±0.05 Hz to 11.31±0.05 Hz in CC and from 11.33±0.04 Hz to 11.31±0.04 Hz in AA, *p*=0.005, Table S14).

We also detected a night by genotype interaction for fast spindle amplitude (F(1, 1356.05)=10.24, *p*=0.001, Table S15), with differences in estimated marginal means shown in Figure 4F: amplitudes increased from night 1 to night 2, but again only in the CC group participants (from 31.8±2.0μV to 32.6±2.0μV, *p*=0.007, Table S16). Fast spindle frequency did not vary across nights or genotype, but fast spindle duration showed a similar pattern to fast spindle amplitude: a genotype by night interaction (F(1, 1356.53)= 7.24, *p*= 0.0072), driven by shorter spindles in CC group during night 1 (795±8 ms vs. 819±8 ms in AA, *p*= 0.02, Table S16) and an increase in spindle length from night 1 to night 2 only in the CC group (from 795±8 ms to 800±8 ms, *p=* 0.03, Table S16). We found no strong evidence for an effect of genotype or night on slow or fast spindle density (Table S13, 15). These fast spindle event-based analyses were supported by spectral analyses of fast sigma power, which showed differential patterns of experience-dependent changes between genotypes (Tables S7, S8).

To summarize these NREM EEG event analyses, only the CC genotype group showed slow-wave and spindle properties – particularly event amplitudes – that were sensitive to experience, sustaining increases on night 2 (post-MST learning) relative to night 1 (baseline). It is possible, then, that attenuated experience-dependent changes in thalamocortical activity contributed to more variable MST performance in the AA group. However, since recent work has highlighted the importance of temporal interrelationships between these thalamocortical oscillations for sleep-dependent memory consolidation (42,55–57), we next tested whether slow-wave coordination also varied across nights and participants.

### Slow wave mediated cortical connectivity during NREM sleep

We analyzed slow wave synchronization during NREM sleep for both genotype groups and nights using multi-taper spectral coherence. Figure 5A illustrates all EEG electrode positions and Figure 5B shows raw EEG traces surrounding a single SW event detected at electrode Fz. To illustrate SW-associated temporal covariance in frontal and occipital EEG, we used Fz SW events (trough times) as triggers to extract +/-2s windows of EEG surrounding each event across both channels, averaging across all windows for each recording night. Figure 5C shows Fz SW event triggered averages at Fz and O1 from one participant of the CC group. Here we can see highly stereotypical SW events detected at Fz (with low variance) during both recording nights, but different average waveforms at O1. During night 1, Fz SW coincide with highly variable activity at O1, where a SW-like waveform is hardly separated from surrounding background activity (Figure 5, C1); in contrast, during night 2 a distinct average SW waveform coordinated with Fz, manifests at O1 (Figure 5, C2).

**Figure 5:**
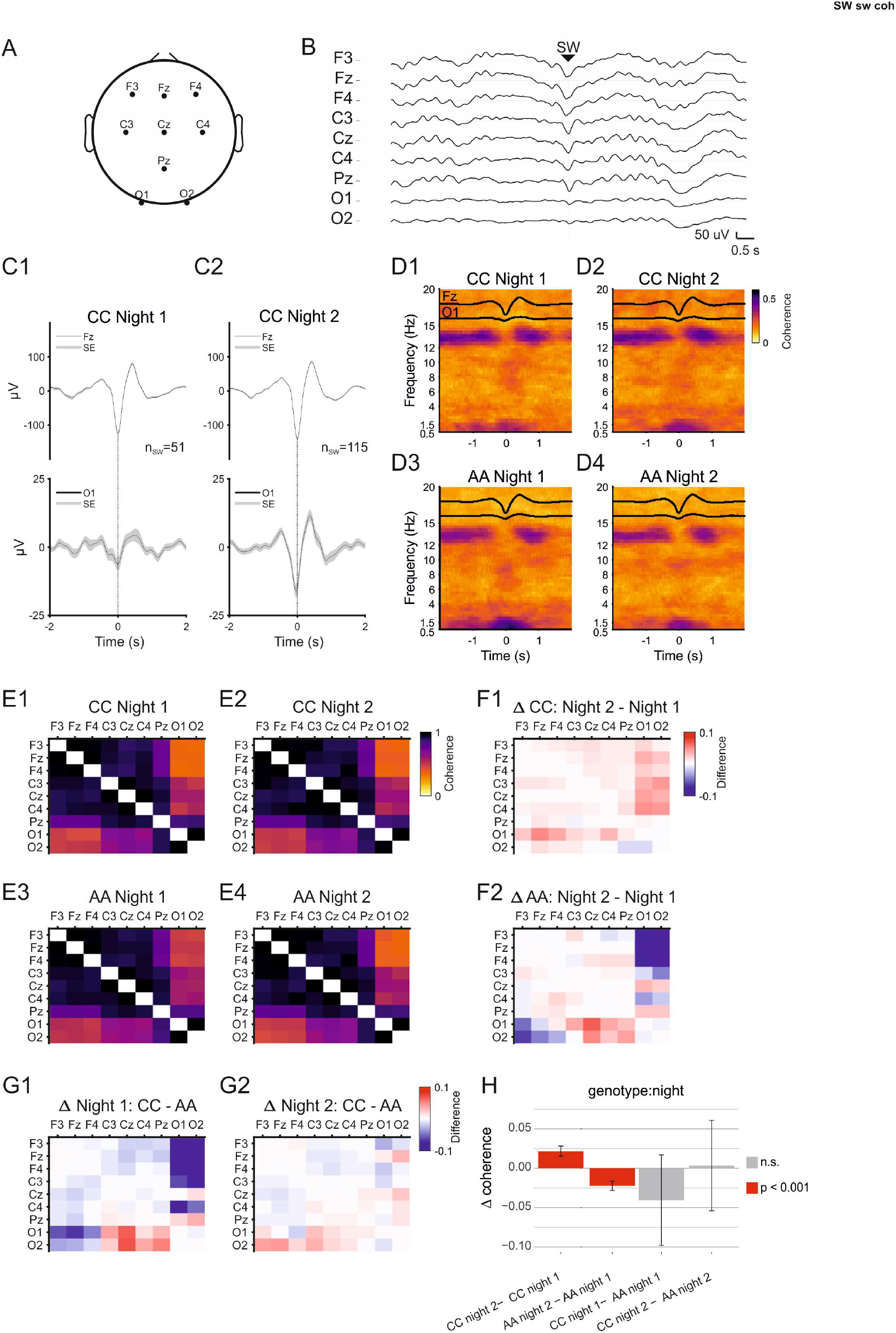
Slow wave triggered coherence. A) A head diagram showing sleep EEG electrode positions. B) Raw data example showing all EEG traces surrounding a typical detected SW event at Fz. C) SW triggered averages from one participant from band pass filtered (0.5-4 Hz) and EEG traces. Individual SW event traces were averaged in windows of +/-2 seconds with the SW trough time set t=0. C1) SW triggered average at Fz and O1 for 51 SW events detected during night 1 N2 sleep in one participant (CC, Night 1), C2) SW triggered averages as described in C1 using, 115 SW events during night 2 from the same participant (‘CC’, Night 2) D) Average SW triggered coherograms. SW triggered data windows (SW trough time at Fz, t = 0) from the seed electrode (Fz) and target electrode (O1) were used to calculate multitaper coherograms for each data window pair. Coherograms were averaged for each participant and then averaged for each group and recording night. In the coherogram darker colors indicate higher coherence. Overlaid black traces are SW wave triggered averages for seed and target electrode (Fz, top and O1, bottom). D1) Average coherogram for CC night1, D2) Average coherogram for CC night 2, D3) Average coherogram for AA night 1, D4) Average coherogram for AA night 2. E) Average SW-triggered coherence values (0.5-1.5 Hz, −0.5 −0.5 s) for all electrode pairs for each genotype group during N2 sleep for both recording nights. E1) CC, during (baseline) night 1; E2) CC, during (learning) night 2, E3) AA, during (baseline) night 1, E4 AA, during (learning) night 2. F) Differences in SW coherence between nights in each genotype group CC (F1), and AA (F2). G) Differences between genotype groups on each night, night 1: CC-AA (G1), night 2: CC-AA (G2). H) Estimated marginal means differences for the factors genotype and night estimated from a linear mixed model analysis of SW event coherence. CC individuals show an increase in SW coherence during night 2, but AA’s show an overall decrease in coherence. Error bars indicate 95% confidence intervals.

Figure 5, panels D1-4 show group-averaged coherograms for the electrode pair Fz-O1 for both recording nights and genotype groups: the most coherent frequency ranges are 0.5-1.5 Hz (SW) and fast spindle coherence (12-15 Hz). We used the average SW coherence (0.5-1.5Hz) during 1s windows surrounding each SW for each electrode pair to construct a cortex-wide SW connectivity matrix. Figure 5, E1-4 show matrices of group averaged coherence values for both genotypes and both recording nights during N3 sleep. All matrices show a gradient of coherence, with highest values between frontal and central electrodes and lowest values between the most distant pairs, i.e. frontal and occipital electrodes.

We calculated the difference between all coherence values and plotted them in the same matrix layout to illustrate differences in slow-wave coherence between nights (Figure 5, F1-2) and genotypes (Figure 5, G1-2). Consistent differences in slow-wave coherence are apparent between recording nights. Slow wave coherence increases in those in the CC group from night 1 (baseline) to night 2 (learning, Figure 5, F1) - but a decrease in the AA group can be seen in the difference matrices for this genotype group (Figure 5, F2). Slow-wave coherence is higher in the AA group on night 1 (mainly frontal to occipital coupling between electrodes F3, Fz, F4 to O1, O2), compared to the CC group (Figure 5, G1). Furthermore, a linear mixed model with subsequent stepwise reduction revealed a genotype by night interaction, indicating a differential effect of learning on slow-wave coherence in CC vs. AA genotypes (genotype x night: F(1, 11182) = 97.37, p<2e-16, Table S17). Both genotypes show changes in slow-wave coherence upon motor learning but a least squares estimation of group marginal means reveals that those in the CC group show a post-learning increase in slow-wave coherence (CC night 1 0.85 ± 0.021, CC night 2 0.87±0.019, p<0.001, Table S18), whereas the AA group show a decrease in overall slow-wave coherence (genotype by night: AA night 1 0.89±0.019 - AA night 2 0.86±0.021, p<0.001, Table S18) after learning. These results are confirmed when using SW coherence of whole noise free NREM sleep epochs (see Supplemental Results and Tables S19, S20). Thus, the CC group showed the increased SW coherence predicted by previous studies (43,58), whereas SW coordination was attenuated following learning in the AA participants.

## DISCUSSION

We performed a recall-by-genotype study (45) to investigate the potential contributions of an SZ-associated SNP, rs1344706, to sleep-dependent memory processing and sleep neurophysiology in healthy volunteers. In summary, 1) all participants showed normal wake/sleep rhythms and there was no evidence for an effect of genotype on any variable describing macro sleep architecture; 2) we observed greater variance in learning and sleep-dependent memory consolidation following a motor task in AA participants homozygous for the variant associated with increased SZ liability; 3) we detected genotype- and learning-dependent effects on SW and fast spindle amplitudes, with the AA group showing attenuated changes in SW and spindle amplitudes after learning; 4) the AA group failed to exhibit the learning-dependent increase in SW coherence evident in the CC genotype group.

### Behavior

Using a motor learning task previously shown to be impaired in SZ patients, we did find evidence for greater variability in overnight improvement and other variables derived from MST in those with the AA genotype at rs1344706, suggesting that rs1344706 may associate with subtle changes motor learning and consolidation.

Motor learning (59) and its sleep-dependent memory consolidation are impaired in patients diagnosed with SZ (60). The key brain areas involved in motor learning and its sleep dependent consolidation, including the neocortex, striatum, thalamus, hippocampus and cerebellum (61,61–64), have all been implicated in the etiology of SZ (65–67). *ZNF804A* has been shown to be highly expressed in these brain regions, particularly the thalamus, hippocampus and cortex (68), hence altered *ZNF804A* function may contribute to changes in brain development and plasticity that influence motor learning and its consolidation (69). Previous studies have shown that variability between individuals during early phases of learning a motor task is higher in patients diagnosed with SZ compared to healthy controls (70), potentially reflecting higher variability in brain anatomy or functional connectivity patterns (71). Whether this variability and the associations of *ZNF804A* derive from neurodevelopmental effects or altered adult neural plasticity remains an open question.

### NREM sleep and neurophysiology

Although we did not observe consistent evidence for an effect of genotype on diurnal rhythmicity or sleep architecture, detailed analyses of overnight EEG did unveil relationships between rs1344706, corticothalamic activity during NREM sleep and neural correlates of motor memory consolidation. We observed several interaction effects between genotype and recording night, where NREM sleep activity appears to be differentially affected by the acquisition of a motor task in AA and CC participants.

#### Spindle oscillations

Previous studies have shown that sleep dependent motor memory consolidation correlates with spindle oscillations (56,72–75). Indeed, a substantial body of work has demonstrated correlations between N2 sleep or spindles with motor memory in healthy participants (76–79), although contradictory studies do exist (80,81). In particular, the individual contributions of slow and fast spindles to memory consolidation are still debated.

We found some evidence supporting a role for slow spindle oscillations (9-12 Hz) in motor memory consolidation, since slow spindle amplitudes appeared to be increased in CC genotype participants during the learning night, whilst decreasing in the AA group. However, we found no evidence of an effect of genotype or night on either slow or fast spindle density. Despite fast spindle density being reduced in first episode (34,82,83) and chronically ill patients (32,33,39) and their first-degree relatives (29,34,82), and the association of spindle properties with polygenic risk scores for SZ (84), our results imply that rs1344706 alone does not impact directly on fast spindle density. Given established polygenic effects in SZ, multiple genetic variants and their interactions are likely to impact cortico-thalamic circuit development. In particular, SNPs linked to ion channel genes like *CACAN1C* (84) are likely to interact with other SNPs to impact corticothalamic development and maturation which might have causal effects on cortico-thalamic oscillatory signatures or NREM sleep (40).

#### Slow oscillations

On average, slow-wave amplitudes increased during the sleep after learning only in the CC genotype group. In addition, SW coherence appears to be differentially modulated after learning between the genotype groups: those in the CC group show an increase in slow-wave coherence, but participants with the AA genotype show a decrease in slow wave coherence during the night that followed motor learning. Previous studies have shown that during early sleep, after motor learning, slow-wave event amplitudes are locally increased in central and parietal areas (85). Slow wave coherence has also been shown to increase during sleep after a declarative memory task (58), and our recent work has demonstrated slow wave coherence increased after motor learning in a control group, but not in patients diagnosed with SZ (43). Our results in individuals homozygous for the ‘A’ allele at rs1344706 seem to be line with these findings and provide a genetic correlate for slow wave phenotypes related to psychosis and SZ.

Recent studies on the rodent homologue of *ZNF804A* suggest it has both a role during development and in adult plasticity (69,86). Our own work in a rodent neurodevelopmental model of SZ has demonstrated that interference in cortico-thalamic development causes severe disruption of slow wave coordination between remote cortical areas and simultaneous desynchronization of spindle and hippocampal ripple oscillations (51). Given the suggested role of *ZNF804A* in cortical and thalamic development we speculate that rs1344706 may have a role in corticothalamic development which itself would be related to impaired coordination of SW activity during sleep. These deep characterizations of genotypic association motivate future mechanistic studies in animal models that enable high-resolution phenotyping corticothalamic circuit development and plasticity.

### Limitations

Currently, we are only beginning to understand complex genotype-phenotype relationships related to SZ. Quantifying brain activity and function directly related to identified neuronal circuits in combination with a Recall-by-Genotype approach is critically advancing our understanding of underlying biological mechanisms. Here we characterized the impact of the rs1344706 variant on sleep neurophysiology but have not directly quantified the effect of the variant on *ZNF804A* gene expression and/or function. Indeed, no consensus has been reached on the function of *ZNF804A* itself yet, although a recent study suggests that *ZNF804A* may regulate RNA synthesis (89). The rs1344706 variant may then reduce the expression of a *ZNF804A* splice variant during prenatal development (87) leading to reduced spine density in cortical excitatory neurons (69,88) - although the exact impact of rs1344706 on corticothalamic development remains to be investigated. Moreover, recent studies show that the impact of rs1344706 on brain function may depend on the carriage of other SZ-associated loci (e.g. in genes *CACNA1C* (90) and *COMT* (91,92)) and environmental risk factors such as cannabis use (93). Thus, the relationship between rs1344706 and its downstream consequences is not easily determined. Furthermore, whilst the intention here was to understand the role of *ZNF804A* in the absence of disease, further work is needed to show that the phenotypes we observed here, are indeed relevant to SZ.

Finally, the small sample size and lack of replication within this study (due to the labor and cost intensive nature of sleep laboratory experiments) naturally limit the strength of our conclusions. Future work will look to improve the efficiency and reach of genetic sleep studies by developing protocols that implement wearable technology to monitor sleep neurophysiology at-home, over extended periods of time and in much larger samples.

## Conclusions

Given the complex network of events linking genetics to brain-wide connectivity and function, how can we best map genomic information to a neurobiological understanding of SZ? Here we show that sleep neurophysiology presents a uniquely powerful opportunity to bridge different levels of analysis: relating genotype to sleep-dependent physiology and environmental factors such as learning, constitutes a rational, neurobiologically-informed approach to delineating causal mechanisms of thalamocortical circuit dysfunction (44). Future translational studies should investigate the influence of *ZNF804A* on slow wave and spindle properties and their coordination in genetic rodent models and patient populations to further elucidate genetic and circuit mechanisms of psychosis and their impacts on sleep, cognition and novel therapies.

## ACKNOWLEDGEMENTS

We are extremely grateful to all the families who took part in the ALSPAC study, the midwives for their help in recruiting them, and the whole ALSPAC team, which includes interviewers, computer and laboratory technicians, clerical workers, research scientists, volunteers, managers, receptionists and nurses. The UK Medical Research Council (MRC) and Wellcome (Grant ref: 217065/Z/19/Z) and the University of Bristol provide core support for ALSPAC. A comprehensive list of grants funding is available on the ALSPAC website (http://www.bristol.ac.uk/alspac/external/documents/grant-acknowledgements.pdf).

Specifically, GWAS data was generated by Sample Logistics and Genotyping Facilities at Wellcome Sanger Institute and LabCorp (Laboratory Corporation of America) using support from 23andMe; PLIKS data at age 18 was funded by MRC grant G0701503/85179 (PI: Stan Zammit); WASI at age 15 was funded by Wellcome Trust and MRC grant 076467/Z/05/Z (PI: George Davey Smith). This research was specifically funded by Elizabeth Blackwell Institute Catalyst Fund grant (ref: 48474) and the MRC Integrative Epidemiology Unit (IEU) [MC_UU_12013/3]. This publication is the work of the authors and NJ Timpson and MW Jones will serve as guarantors for the contents of this paper.

NJT is a Wellcome Trust Investigator [202802/Z/16/Z to NJT] and works within the University of Bristol National Institute for Health Research (NIHR) Biomedical Research Centre (BRC). LJC is supported by NJT’s Wellcome Trust Investigator grant [202802/Z/16/Z]. NJT and LJC work in the MRC Integrative Epidemiology Unit (IEU) at the University of Bristol which is supported by the MRC [MC_UU_00011] and the University of Bristol. NJT is supported by the Cancer Research UK (CRUK) Integrative Cancer Epidemiology Programme [C18281/A19169]. CH was an academic foundation doctor supported by North Bristol NHS Trust during her time on the project. KE was funded through a PhD studentship at the MRC Integrative Epidemiology Unit. MT’s time on the project was funded through an Elizabeth Blackwell Institute postgraduate fellowship. CD’s time on the project was funded through the Elizabeth Blackwell Institute Catalyst Fund grant.

MWJ was awarded an MRC Senior Non-Clinical Research Fellowship (G1002064) that part-funded this work. UB received a Lilly Innovation Fellowship Award through Eli Lilly & Co Ltd UK.

