## Supplemental Information for "Schizophrenia-associated variation at *ZNF804A* correlates with altered experience-dependent dynamics of sleep slow-waves and spindles in healthy young adults"

### Table of Contents


**SUPPLEMENTAL METHODS** 3
**Avon Longitudinal Study of Parents and Children (ALSPAC) cohort description** 3
**Participants** 5
**Supplemental Procedures** 7
**Behavioral Analyses** 8
**Polysomnography Analyses** 11
**Statistical Methods Overview** 13
**SUPPLEMENTAL RESULTS** 15
**Recruitment** 15
**Motor Sequence Task performance** 15
**Diurnal rhythmicity, subjective and objective sleep quality** 16
**Spectral analysis of whole NREM epochs** 17
**NREM event detection** 18
**Coherence during whole NREM epochs** 19
**SUPPLEMENTAL DISCUSSION** 20
**NREM power** 20
**NREM coherence** 21
**SUPPLEMENTAL TABLES** 23
**Table S1 - Overview of statistical analysis of non-EEG data** 23
**Table S2 - Overview of statistical analysis of sleep EEG data** 24
**Table S3 - Characteristics of ALSPAC participants recruited and not recruited to the study** 25
**Table S4 - Habitual sleep behavior and diurnal rhythms derived from actigraphy** 26
**Table S5 - PSG-derived sleep architecture across groups and sessions** 27
**Table S6 - Sleep architecture in participants with PSQI scores above and below 5** 28
**Table S7 - Linear mixed model results for NREM power** 29
**Table S8 - Estimated marginal means of NREM power across all electrodes** 30
**Table S9 - Linear mixed model results for NREM SW event properties** 31
**Table S10 - Estimated marginal means of NREM SW event properties across all electrodes:** 32
**Table S11 - Linear mixed model results for NREM delta wave event properties** 33
**Table S12 - Estimated marginal means of NREM delta wave event properties across all electrodes** 34
**Table S13 - Linear mixed model results for NREM slow spindle event properties** 35
**Table S14 - Estimated marginal means of NREM slow spindle event properties across all electrodes** 36
**Table S15 - Linear mixed model results for NREM fast spindle event properties** 37
**Table S16 - Estimated marginal means of NREM fast spindle event properties across all electrodes** 38
**Table S17 - Linear mixed model results for SW triggered slow coherence** 39
**Table S18 - Estimated marginal means of SW triggered SW coherence across all electrodes** 39
**Table S19 - Linear mixed model results for whole epoch NREM coherence** 40
**Table S20 - Estimated marginal means of whole epoch NREM coherence across all electrodes** 41
**SUPPLEMENTAL FIGURES** 42
**Figure S1 - CONSORT flow diagram of participant recruitment and exclusions prior to analysis** 42
**SUPPLEMENTAL REFERENCES** 43

### Supplemental Methods

#### Avon Longitudinal Study of Parents and Children (ALSPAC) cohort description

*Description of study numbers*

Pregnant women resident in Avon, UK with expected dates of delivery 1st April 1991 to 31st December 1992 were invited to take part in the study. The initial number of pregnancies enrolled is 14,541 (for these at least one questionnaire has been returned or a "Children in Focus" clinic had been attended by 19/07/99). Of these initial pregnancies, there was a total of 14,676 fetuses, resulting in 14,062 live births and 13,988 children who were alive at 1 year of age. When the oldest children were approximately 7 years of age, an attempt was made to bolster the initial sample with eligible cases who had failed to join the study originally. As a result, when considering variables collected from the age of seven onwards (and potentially abstracted from obstetric notes) there are data available for more than the 14,541 pregnancies mentioned above. The number of new pregnancies not in the initial sample (known as Phase I enrolment) that are currently represented on the built files and reflecting enrolment status at the age of 24 is 913 (456, 262 and 195 recruited during Phases II, III and IV respectively), resulting in an additional 913 children being enrolled. The phases of enrolment are described in more detail in the cohort profile paper and its update. The total sample size for analyses using any data collected after the age of seven is therefore 15,454 pregnancies, resulting in 15,589 fetuses. Of these 14,901 were alive at 1 year of age. A 10% sample of the ALSPAC cohort, known as the Children in Focus (CiF) group, attended clinics at the University of Bristol at various time intervals between 4 to 61 months of age. The CiF group were chosen at random from the last 6 months of ALSPAC births (1432 families attended at least one clinic). Excluded were those mothers who had moved out of the area or were lost to follow-up, and those partaking in another study of infant development in Avon.

*Genotyping description*

ALSPAC children were genotyped using the Illumina HumanHap550 quad chip genotyping platforms by 23andme subcontracting the Wellcome Trust Sanger Institute, Cambridge, UK and the Laboratory Corporation of America, Burlington, NC, US. The resulting raw genome-wide data were subjected to standard quality control methods. Individuals were excluded on the basis of gender mismatches; minimal or excessive heterozygosity; disproportionate levels of individual missingness (>3%) and insufficient sample replication (IBD < 0.8). Population stratification was assessed by multidimensional scaling analysis and compared with Hapmap II (release 22) European descent (CEU), Han Chinese, Japanese and Yoruba reference populations; all individuals with non-European ancestry were removed. SNPs with a minor allele frequency of < 1%, a call rate of < 95% or evidence for violations of Hardy-Weinberg equilibrium (P < 5E-7) were removed. Cryptic relatedness was measured as proportion of identity by descent (IBD > 0.1). Related subjects that passed all other quality control thresholds were retained during subsequent phasing and imputation. 9,115 subjects and 500,527 SNPs passed these quality control filters.

ALSPAC mothers were genotyped using the Illumina human 660W-quad array at Centre National de Génotypage (CNG) and genotypes were called with Illumina GenomeStudio. PLINK (v1.07) (1) was used to carry out quality control measures on an initial set of 10,015 subjects and 557,124 directly genotyped SNPs. SNPs were removed if they displayed more than 5% missingness or a Hardy-Weinberg equilibrium P-value of < 1E-6. Additionally, SNPs with a minor allele frequency of less than 1% were removed. Samples were excluded if they displayed more than 5% missingness, had indeterminate X chromosome heterozygosity or extreme autosomal heterozygosity. Samples showing evidence of population stratification were identified by multidimensional scaling of genome-wide identity by state pairwise distances using the four HapMap populations as a reference, and then excluded. Cryptic relatedness was assessed using an IBD estimate of more than 0.125 which is expected to correspond to roughly 12.5% alleles shared IBD or a relatedness at the first cousin level. Related subjects that passed all other quality control thresholds were retained during subsequent phasing and imputation. 9,048 subjects and 526,688 SNPs passed these quality control filters.

*Imputation description*

477,482 SNP genotypes in common between the sample of mothers and sample of children were combined. SNPs with genotype missingness above 1% due to poor quality were removed (11,396 SNPs removed). 321 subjects were removed due to potential ID mismatches. This resulted in a dataset of 17,842 subjects containing 6,305 duos and 465,740 SNPs (112 were removed during liftover and 234 were out of HWE after combination). Haplotypes were estimated using ShapeIT (v2.r644) which utilizes relatedness during phasing. A phased version of the 1000 genomes reference panel (Phase 1, Version 3) was obtained from the Impute2 reference data repository (phased using ShapeIt v2.r644, haplotype release date Dec 2013). Imputation of the target data was performed using IMPUTE V2.2.2 (2,3) against the reference panel (all polymorphic SNPs excluding singletons), using all 2,186 reference haplotypes (including non-Europeans). This gave 17,842 mothers and children eligible for study with available genotype data. Subsequent consent withdrawals have left 17,825 individuals for study.

*Ethical approval*

ALSPAC has its own Ethics and Law Committee that reviews all proposals for new data collection and approves policies for data handling and analysis. Ethical approval for the study was obtained from the ALSPAC Ethics and Law Committee and the Local Research Ethics Committees. Proposals for new data collection are also approved by the Local Research Ethics Committees (LRECs). Consent for biological samples has been collected in accordance with the Human Tissue Act (2004). Informed consent for the use of data collected via questionnaires and clinics was obtained from participants following the recommendations of the ALSPAC Ethics and Law Committee at the time.

*Study data*

The ALSPAC study website contains details of all the data that is available through a fully searchable data dictionary (http://www.bristol.ac.uk/alspac/researchers/our-data/).

#### Participants

rs1344706 is located on chromosome 2 at position 185,778,428 bp (genome build GRCh37). In the ALSPAC genetics data described above, the minor allele (C) occurs at a frequency of 40.0% (as compared to 37.8% in the European arm of the 1000Genomes dataset) and the variant is imputed with an information score of 0.995. Each individuals genotype at rs1344706 was determined based on genotype probabilities in the imputed data, such that the most likely genotype was assigned. Male participants of European ancestry were then selected for invite based on their being homozygous either for the rs1344706 allele associated with increased liability for schizophrenia (AA group) or for the alternative allele (CC group), with an equal number of invites being sent to AA and CC individuals. We restricted our study to males due to the evidence for heterogeneity of SZ by sex and the higher rates of disease observed in males (4). Following a positive response from a participant, telephone screening established eligibility prior to sleep clinic visits. Eligible participants were: (1) aged 20 years or over; (2) male; (3) non-smokers; (4) of European ancestry; (5) in good physical and mental health with no history of diagnosed sleep disorders; (6) able to give informed consent as judged by the investigator. Participants were excluded if: (1) they had current substance dependence (other than caffeine); (2) they had a substantive current or past illness; (3) were taking any medications that may affect or induce sleep; (4) worked at night. Participant eligibility was then verified on arrival at the sleep clinic through further standardized screening questions and completion of the Bristol Sleep Profile, BSP, (5) and The Pittsburgh Sleep Quality Index, PSQI, (6). A CONSORT flow diagram detailing the recruitment process is shown in Figure S1. Two rounds of data collection were carried out with 26 participants recruited in the first round and 21 in the second round; of these, 46 individuals (25 AA and 21 CC) completed the study. At no point were participants made aware of their genotype status. We checked for an association between genotype group and several potential confounding factors: maternal education (a binary trait to indicate which mothers had A-Level or degree level qualifications); maternal social class (a six category definition based on mother's occupation and ranging from professional to unskilled); psychosis-like symptoms (PLIKS) at age 18 (a binary trait to indicate the presence of one or more symptoms based on data from a face to face interview to determine psychosis like symptoms); intelligence at age 8 (based on the Wechsler Intelligence Scale for Children (WISC); and intelligence at age 15 (based on the Wechsler Abbreviated Scale of Intelligence). In these analyses, a Pearson Chi-square test was used for categorical variables and a Wilcoxon rank-sum (Mann-Whitney) test for continuous variables. We used the same method to check for differences in potential confounding factors between those invited to the study and those who were recruited.

#### Supplemental Procedures

*Clinic routine*

The data collection phase for each participant was approximately two weeks long, beginning and ending with a night spent in a sleep suite at the Clinical Research and Imaging Centre at the University of Bristol during which a polysomnography (PSG) was performed (Figure 1). On the first visit, participants completed Pittsburgh Sleep Quality Index (PSQI) (6) and Bristol Sleep Profile (BSP) (5) which were used to assess self-rated sleep quality and to indicate any specific sleep disturbance, respectively. Each participant also completed the Edinburgh Handedness Inventory (7) to ascertain participant handedness ahead of the motor sequence task (MST) (see below). Participants were issued with an actigraphy watch (MotionWatch 8, CamNtech, UK) to wear until study completion and a diary in which to record information about their bedtime, wake/rise times, naps, daytime activities and caffeine and alcohol intake.

Each participant stayed in the sleep suite for two nights, one baseline recording night (night 1) and a second night that included the MST (night 2), with approximately two weeks between visits. Following PSG electrode placement and bio-calibration, participants followed their usual evening routine and were encouraged to go to bed at their usual bedtime. In the morning, participants were woken as close as possible to their usual wake time. After each PSG recording, participants completed the St Mary's Hospital Sleep Questionnaire (SMH) (8) and the Leeds Sleep Evaluation Questionnaire (LSEQ) (9) to collect information about subjective experience of their night in the sleep laboratory. On the second night, participants were trained on the MST (see below for details) two hours before their planned bedtime. They were then tested on the MST after electrode removal the following morning.

*Motor Sequence Task*

We used a motor sequence task (MST) to test sleep dependent consolidation of motor memory (10–12) implemented in the Matlab environment using psychtoolbox (13), kindly donated by Dara Manoach (Harvard Medical School, Boston, MA). On their second visit participants performed the MST approximately two hours before they went to bed (training) and again the following morning (test). During the MST, participants were asked to press four numerically labeled keys on a computer keypad in a five-element sequence (4-1-3-2-4) with the fingers of their non-dominant hand, repeating “as quickly and accurately as possible” for 30 seconds. The numeric sequence was visible throughout the trial and dots underneath provided visual feedback for each keystroke. During both training and test sessions, participants alternated tapping and resting for 30 seconds for a total of 12 trials. Prior to completing the MST, the participant completed the Stanford Sleepiness Scale (14) to quantify vigilance levels.

*Actigraphy*

The MotionWatch 8 (CamNtech, UK) is a wrist worn activity monitor that contains a miniature accelerometer to allow measurement and recording of physical movement of the wrist, providing a close correlation to whole body movement. Participants were asked to wear the ‘actiwatch’ for the entire period between their clinic visits, removing it only during water-based activities (e.g. swimming, bathing) and those activities which might result in the actiwatch being damaged (e.g. rugby).

*Polysomnography (PSG)*

A standard in-laboratory, overnight PSG, including video and audio recording, was performed using Embla® N7000 PSG amplifier and RemLogic software (Natus Medical Inc., California). Nine electrodes were placed according to the 10-20 system (at F3, Fz, F4, C3, Cz, C4, Pz, O1 and O2) and data acquired using Cz as reference and a standard PSG recording montage. Additional electrodes were placed to monitor eye movements (electrooculography), submental muscle activity (electromyography) and heart rate (electrocardiogram) throughout the recording.

#### Behavioral Analyses

All behavioral and questionnaire data was analyzed using Stata v14.2 (15) unless stated otherwise. An overview of the non-EEG analyses performed is shown in Table S1.

*Questionnaires*

All paper questionnaires (Pittsburgh Sleep Quality Index, Bristol Sleep Profile, Edinburgh Handedness Inventory, Stanford Sleepiness Scale) were manually scored and transcribed to Excel sheets. A second researcher checked at least 25% of the data entries per questionnaire, with additional checks being carried out if errors were identified.

*Motor Sequence Task*

Results from the Stanford Sleepiness Scale, applied to assess self-reported vigilance at the time of the MST, were compared across genotype groups using a two-sample two-sided t-test (with unequal variances). The primary outcome measures from the MST were (1) the number of correct sequences per 30 second epoch, which reflects both the speed and accuracy of performance; and (2) button press latency (reaction time, RT) in milliseconds (ms) during correct sequences. These measures were derived for each 30-second trial in the evening (training) and morning (test) sessions. For each outcome measure, 'training performance' was defined as the average of the last three training trials and 'test performance' was defined as the average of the first three test trials. Overnight improvement was calculated as the percentage change in each outcome measure from training performance to test performance (16).

Participant performance in the MST was initially compared across genotype groups (AA versus CC) by plotting average learning curves. Sleep dependent memory processing of motor learning was then formally compared across genotype groups by two approaches. Firstly, the mean and variance of overnight improvement measures were compared using two-sample two-sided t-tests (with unequal variances) and two-sample variance comparisons, respectively. Secondly, a linear mixed model framework was applied where training (evening) and test (morning) performance were considered repeat observations. The regression was fitted via restricted maximum likelihood (REML) using a generalized Satterthwaite approximation to estimate degrees of freedom. Session (training or test) and genotype were modelled as fixed effects, whereas participant identity was modelled as a random effect. Interactions between fixed effects were added to the final model if a likelihood ratio test comparing nested models with and without the interaction parameter suggested an improvement to model fit ($p<0.05$, maximum likelihood models, ML). The assumptions of the linear regression model were checked by plotting histograms and Q-Q plots of residuals from the models. In addition, a Levene's robust test for equality of variance across groups (within session) was applied (17)

*Actigraphy*

The habitual sleep behavior of participants was evaluated using actigraphy data collected in the two weeks between clinic visits. Actigraphy data was manually annotated in

MotionWare (CamNtech, UK) using information from participant diaries and participant submitted event markers to indicate sleep and rise times. Periods for which the participant had removed the watch were set to missing. An automated scoring algorithm determined 'sleep onset' and 'sleep offset' for each night, except where diary information and/or activity counts contradicted these times. The 'sleep analysis' function in MotionWare was used to derive time in bed (TIB) (total elapsed time between the 'Lights Out' and 'Got Up' times), total sleep time (TST) (the total time spent in sleep according to the epoch-by-epoch wake/sleep categorization), sleep efficiency (TST/TIB) (actual sleep time expressed as a percentage of time in bed), sleep onset latency (SOL) (the time between 'Lights Out' and 'Fell Asleep') and fragmentation index (FI) (the sum of the 'Mobile time (%)' and the 'Immobile bouts <= 1min (%)'), an indication of the degree of fragmentation of the sleep period and an indicator of sleep quality. Measures were then averaged across all available nights for each participant.

We used non-parametric circadian rhythm analysis (NPCRA) to quantify the regularity of daily and weekly sleep wake rhythms (18). A modified version of the algorithm implemented in the MotionWare software was kindly provided by Eus Van Someren. The algorithm was modified such that periods of missing data (where participants had removed their watch) could be excluded from the analysis. Periods for exclusion were defined either by the start and end time of the missing data period (for periods >1 hour and < 3 hours), or by the start time and the start time plus 24 hours (for periods >3 hours). Only participants with at least seven days of data remaining after exclusions were included in the analysis.

The following variables were derived for comparison across genotype groups: (i) Interdaily Stability (IS): the degree of regularity in the activity-rest pattern with a range of 0 to 1 where a value of 0 indicates a total lack of rhythm and a value of 1 indicates a perfectly stable rhythm; (ii) Intra-Daily Variability (IV): the degree of fragmentation of activity-rest periods with a theoretical range of 0 to 2 with higher values indicating higher fragmentation; (iii) Relative Amplitude (RA): the difference between the highest and lowest activity levels and has a range of 0 to 1 with higher values indicating a rhythm with higher amplitude; (iv) Least 5 Average (L5): the average activity level for the sequence of the least five active hours; and (v) Most 10 Average (M10): the average activity level for the sequence of the highest (most) ten active hours. Sleep architecture and NPCRA measures derived from the actigraphy data were compared across genotype groups using a Wilcoxon rank-sum (Mann-Whitney) test.

#### Polysomnography Analyses

*Sleep Architecture*

PSG data were manually scored by an experienced expert (blinded to participant genotype) based on AASM criteria (19) using REMLogic software (Natus Europe GmbH, Germany). Each 30-second epoch was visually classified into stages (Wake, NREM1, 2, 3 and rapid-eye movement). Awakenings were scored when one or more 30-second epoch was classified as wake following initial sleep onset. Individual sleep continuity and architecture was quantified using standard variables: time in bed (TIB), total sleep time (TST), sleep latency (SOL), wake after sleep onset (WASO) and sleep efficiency. Sleep stages are presented as the percentage of TST. These outcome measures were compared across genotype groups and nights using the same linear mixed model framework as used to assess MST performance (fitted in Stata 14.2 (15) using REML). Participant identity was fitted as a random effect and genotype group and recording night (night 1: baseline, night 2: learning) were fitted as fixed effects. The presence of interactions between fixed effects were evaluated via a likelihood ratio test comparing nested models with and without the interaction parameter.

*Spectral and Coherence Analysis of NREM Sleep Epochs*

Sleep scored and epoched EEG data were manually reviewed (noisy epochs and channels were removed). We then performed spectral analysis of all channels of whole epochs of sleep stages N2 and N3 via multi-tapered spectra (Tables S7, S8) and coherence (Tables S19, S20) using the Chronux toolbox (Mitra and Bokil, 2008, www.chronux.org) using 9 tapers, 10s window, 1s sliding window.

*NREM Event Detection (slow-wave, delta, spindle)*

We focused further EEG analyses on NREM sleep for three primary reasons: (1) NREM neurophysiology encompasses hallmark thalamocortical oscillations – slow-waves and spindles – that can be readily identified in scalp EEG; (2) sleep-dependent memory in the MST has been shown to correlate with NREM neurophysiology in healthy adults (21–23); and (3) there is evidence for altered NREM sleep neurophysiology in SZ (12,24–29).

NREM sleep is subdivided into stages N2 and N3. N2 is rich in spindles, which reflect rhythmic activity in thalamocortical circuits and can be further classified as slow (9-12 Hz) spindles with a fronto-central focus, or fast (12-16 Hz) spindles with a centro-parietal focus (30–32). N3 reflects deeper sleep and features the densest and highest-amplitude slow (<1.5 Hz) and delta waves (>1.5-4 Hz intrinsic frequency), which signify the coordinated firing of neocortical neurons and appear most prominently on electrodes over frontal cortices (33–37).

We automatically detected slow wave (SW), delta wave and spindle events using custom Matlab algorithms as described previously (29,38) (code is available at https://gitlab.com/ubartsch/sleepwalker). Slow and delta waves were detected from 0.25-4Hz band pass filtered EEG: the whole EEG trace (visually identified noisy epochs were manually removed) was converted to a z-score and threshold crossings above 3.5 standard deviations (SD) from mean amplitude were detected as candidate events. These candidate events were only accepted as slow-wave if they fell within the following parameter ranges: amplitude 50-300 µV; duration (length in time) 0.2-3s; minimum gap between slow-wave to be considered separate events 0.5s. Slow-wave events are single wave negative threshold crossings with an intrinsic frequency (the inverse of the time difference between first peak and trough multiplied by 2) below 1.5Hz, whereas delta waves are negative single wave threshold crossings with a frequency above 1.5Hz.

To detect spindles, EEG traces were band pass filtered (9–16 Hz), z-scored, rectified and an envelope of the rectified signal was determined using a cubic spline fit to the maxima of the rectified signal. Candidate spindle events were detected as threshold crossings above 3.5 SD of the envelope, then classified as spindles if the absolute amplitude was in the range of 25-500µV; their duration was in the range of 0.25-3 s; and the minimum gap between spindles to be considered separate events 0.25 s; the threshold to detect start and end times for spindle events was set at 1.5 SD. Slow and fast spindles were dissociated based on the average intrinsic oscillation frequency (1/period) with spindles at 9-12Hz labelled as slow spindles and spindles at 12-16 Hz labelled as fast spindles. Following event detection (slow waves, delta waves and spindles), the event density (#/min), frequency (Hz), length (= duration, seconds) and mean peak-to-peak amplitudes (µV) were calculated for each electrode.

To compute average waveforms, we extracted data surrounding detected event times (time of maximum amplitude) and calculated average waveforms by averaging all data in the window (Figure 4). Potential differences between average waveforms at every time bin were assessed in exploratory analyses using a Wilcoxon ranksum test without correction for multiple testing.

*Event triggered coherence analysis*

We further analyzed connectivity using detected SW events as the trigger for a fine-grained analysis of coherence occurring near SW events. An average SW triggered coherogram was calculated using multi-tapered coherograms using the Chronux toolbox (www.chronux.org). SW negative peak times were used as t=0 to collect ±2s of raw EEG around each SW event. The event triggered coherograms were then calculated using 3 tapers, a 1s sliding data window, 50ms steps and were then averaged for each electrode pair (per subject) and then averaged across genotype groups and recording night. The average SW coherence was calculated from the coherograms using a [-0.5 - 0.5 s] and [0.5-1.5 Hz] window, and these average values were then visualized as coherence matrix showing the average coherence values for each genotype group and recording night.

#### Statistical Methods Overview

We used a variety of statistical methods to analyze qualitative, behavioral, actigraphy and EEG data. Statistical methods for the analysis of all behavioral data, including questionnaires, are summarized in Table S1, statistical approaches for the analysis of EEG data is summarized in Table S2.

Boxplots summarize data as median with boxes indicating the $25^{th}$ and $7^{th}$ quartile of data, whiskers extend to 1.5* interquartile range (IQR), i.e. $q_3 + 1.5 \times (q_3 - q_1)$ and $q_1 - 1.5 \times (q_3 - q_1)$ and values outside this range are displayed individually.

NREM event properties, spectral power and coherence measures (derived as described above) were compared across genotype groups, electrodes, recording nights (night 1: baseline, night 2: learning) and sleep stages (N2, N3) using a linear mixed model framework using the lme4 package in R (39). Participant ID was a random effect; genotype, group, electrode, night and sleep stage were fixed effects and an interaction term for genotype * night was included by default. The full model was of the general form *y ~ genotype + night + electrode + sleep_stage + (genotype * night) (1|ID)*, where y is any derived sleep EEG variable. Then a stepwise reduction procedure was employed to remove non-significant terms of linear mixed models, where terms are removed recursively beginning with high level interactions as implemented in the function step of the R package lmerTest (40). Predicted marginal means (designs were not balanced) for each factor and factor interactions were calculated using the R package lsmeans (41). All R analysis routines were implemented using R Studio (rstudio.com) and Microsoft R Open (mran.microsoft.com/open). As in the main txt, results presented herein are mean ± standard error (SE) unless stated otherwise.

### Supplemental Results

#### Recruitment

Data were collected from 47 participants (25 AA and 22 CC). Recruited participants (N=47) did not appear to differ by genotype group or differ from those invited but not recruited (N=430) based on relevant characteristics (Table S3). One CC participant was excluded due to non-completion of the study, data from four participants (2 AA and 2 CC) were excluded due to atypical PSG recordings and a further two participants (1 AA and 1 CC) were excluded based on their failure to perform the MST as instructed (see below for further explanation). We therefore present results for 40 participants (Figure S1).

#### Motor Sequence Task performance

All participants completed evening and morning sessions of the MST; vigilance during the task was assessed using the Stanford Sleepiness Scale and did not differ between genotypes (Table 2). Based on a visual inspection of individual learning curves, data from two participants was deemed to be unreliable on the basis that one had poor performance pre-sleep and a high standard error of the mean (SEM) for reaction time (RT) and the second had a peak in SEM for RT in post-sleep trials 1-3 (i.e. those used to determine test performance). When the performance of these two participants was considered alongside others (within their respective genotype groups), in both cases their improvement metrics (number of correct sequences and RT) fell out with the previously defined limits for outliers (below $Q_1$ – 1.7239*IQR or above $Q_3$ + 1.7239*IQR) indicating unusually poor performance of the task. For this reason, these two participants were excluded from all statistical analyses.

#### Diurnal rhythmicity, subjective and objective sleep quality

Actigraphy data were available for 35 of the 40 participants due to loss of data through technical failures (n=4) and one participant regularly removing his watch overnight. NPCRA analysis was restricted to the 34 participants producing seven or more consecutive days of data after the removal of missing periods. Actigraphy-based analyses of participants' habitual sleep behavior revealed no differences in diurnal sleep routines between genotype groups (Table S4).

Of the 37 (out of 40) participants who correctly completed the PSQI questionnaire, 31 had total scores indicative of good sleep quality (PSQI<=5) and six (two AA and four CC) had scores indicative of poor sleep quality (PSQI>5) (6). Based on polysomnography during clinic nights, participants in both genotype groups showed normal wake/sleep rhythms with total sleep time (TST) and percentage of TST spent in any sleep stage consistent with published ranges (see example hypnograms in Figure 3 A**,** B). For those outcome measures that could be derived from both actigraphy data (during the two-week period between clinic nights) and PSG (during clinic nights), we saw good overall concordance between mean values, showing that clinic visits were representative of 'normal' nights. There was no evidence for an effect of genotype on any variable describing sleep architecture (Figure 3 C-F, Table S5). No difference in sleep quality was indicated in the 6 participants with a PSQI > 5 when measured objectively by actigraphy and PSG (Table S6).

Overall, questionnaire, actigraphy and PSG data show that diurnal rhythms and sleep architecture of AA and CC groups were similar; subsequent analyses are therefore not confounded by frank sleep disruption in either group.

#### Spectral analysis of whole NREM epochs

We further analyzed spectral properties of whole 30 second NREM sleep epochs. We focused on NREM sleep since there is previous evidence for NREM sleep phenotypes in SZ. EEG traces from noise free NREM epochs were analyzed using multitaper spectral analysis (29,42,43).

*Slow Wave power*

Using a linear mixed model approach with subsequent stepwise reduction to remove non-significant terms we found evidence for a night effect on SW power between recording nights 1 and 2 in ($F(1,1373.03) = 7.07$, $p= 0.008$, Table S7). Although there was no main effect of genotype on SW power we detected a significant interaction between genotype and night term ($F(1,1373.03) = 9.13$, $p= 0.003$). Table S8 displays the least squares estimated marginal means for each group and night across all recorded electrodes. We observed an increase in SW power in the CC group (CC night 1: 19.84 ± 0.39 dB, CC night 2: 19.99±0.39 dB, $p= 0.01$) and reduction of SW power during night 2 compared to night 1 in the AA group (AA night 1: 20.68±0.35 dB, AA night 2: 20.45±0.35 dB, $p = 0.008$).

*Delta wave power*

Using a linear mixed model, we obtained similar results to SW power for delta power. We observed both a night effect ($F(1, 1373.02)= 12.89$, $p=0.00034$) and a genotype x night interaction ($F(1,1373.02)= 17.66$, $p= 2.81e-05$, Table S7) in the final stepwise reduced model. Estimated marginal means show an increase in delta power after learning in CC individuals (CC night 1: 11.69 ± 0.35 dB, CC night 2: 11.85±0.35 dB, $p= 0.016$, Table S8) but a decrease in AA individuals (AA night 1: 12.27 ±0.31 dB, AA night 2: 12.05±0.31 dB, $p= 0.0003$, Table S8).

*Slow sigma power*

We analyzed slow sigma power (9-12 Hz) during NREM sleep using the same linear mixed model approach as for slow and delta power. Results for slow sigma power mimic those obtained for delta power, where we see an overall night effect ($F(1, 1373.02)= 6.89$, $p=0.009$) , and a genotype by night interaction ($F(1,1373.02)= 12.50$, $p= 0.0003$, Table S7). When we estimated the marginal means we found that slow sigma power increases in the CC group (CC night 1: 0.38 ± 0.52 dB, CC night 2: 0.60±0.52 dB, $p=0.0134$, Table S8) but decreases in the AA group (AA night 1: 0.96±0.47 dB, AA night 2: 0.75±0.47 dB, p= 0.0088 , Table S8).

*Fast sigma power*

We analyzed multi-taper estimated fast sigma power (12-15Hz) during NREM sleep using the same linear mixed model approach. The final stepwise reduced model shows a clear effect of night on overall sigma power in both genotypes (F(1, 1373.02) = 6.58, p= 0.010, Table S7). We found no evidence for a genotype effect on sigma power, but a genotype x night interaction was included in the final stepwise reduced linear mixed model (F(1, 1373.01) = 7.25, p = 0.007, Table S7). Estimating the predicted marginal means, only those in the AA group show a decrease in fast sigma power after the motor learning during night 2 (AA night 1: -0.77 ±0.47 dB, AA night 2: -0.97 ±0.47 dB, p= 0.010, Table S8) but no change in CC individuals.

In summary, spectral analyses of whole NREM epochs are capable of discerning gene by experience interactions during sleep.

#### NREM event detection

Slow wave and fast spindle event results are described in the main text.

*Delta waves*

We also analyzed delta wave (intrinsic frequency between 1.5 – 4 Hz) event properties using linear mixed models. We observed an effect of night (F(1,1364.03) = 10.96 , p= 0.0001, Table S11) and an interaction effect of night and genotype (F(1,1364.03) = 9.50, p= 0.0021, Table S11) on delta wave amplitude. Estimated marginal means revealed only those in the CC group showed a significant increase in delta amplitude after learning (CC night 1: 92.17 ± 4.04 µV, CC night 2: 95.08± 4.04 µV, p = 9.6e-04, Table S12). We also observed a main effect for night on intrinsic delta frequency (F(1,1365.15)= 6.57, p=0.010, Table S11). Delta waves increase in intrinsic frequency in both genotype groups during the learning night compared to the baseline recording night (Table S12).

*Slow spindles*

When we entered slow spindle (9-12 Hz) properties into linear mixed model analyses, we observed a trend for a night effect (F(1, 1331.05)= 3.70, p= 0.055) and a night by genotype interaction effect (F(1, 1331.06)= 8.42, p= 0.004) on slow spindle amplitude in the step-wise reduced linear mixed model (Table S13). Slow spindle amplitude showed a trend for an increase after leaning in the CC group during night 2 (CC night 1: 31.07 ± 2.02 µV, CC night 2: 31.70 ± 2.02 µV, p=0.06), but decreased in the AA group in comparison to night 1 (AA night 1: 33.80± 1.82 µV, AA night 2: 33.15 ± 1.82 µV, p=0.03) (Table S14). We also observed a night effect on slow spindle frequency (F(1,1332.3) = 9.2693, p=0.002376, Table S13). Both genotype groups show a reduction in slow spindle frequency after motor learning (Table S14).

#### Coherence during whole NREM epochs

We calculated spectral coherence – *i.e*. the covariance of oscillations across slow- and spindle frequencies – for all electrode pairs from whole, noise-free epochs of N2 and N3 sleep. Coherence values were calculated using multitaper coherency estimates based on the hypothesis that coherent phase relationships between oscillating signals recorded across different electrodes reflect functional interactions (Singer, 1999; Greenblatt et al., 2012; Fries, 2015).

*Slow coherence*

We analyzed slow wave coherence in all artefact-free 30 second epochs of N2 and N3 sleep. When we entered all electrode-pair coherence values into a linear mixed model with subsequent stepwise reduction, the final model showed a night effect (F(1, 5554.4) = 5.0623, p=0.025, Table S19) and a genotype x night interaction (F(1, 5554.4) = 24.57, p= 7.37e-07) indicating a differential effect of learning on slow-wave coherence in CC vs. AA genotypes. Least squares estimation of group marginal means showed that those in the CC group show an overall increase in slow-wave coherence (CC night 1: 0.85 ± 0.02, CC night 2: 0.86 ± 0.02, p=0.024, Table S20), whereas AA individuals show a decrease in overall slow-wave coherence (AA night 1: 0.89± 0.02, AA night 2: 0.86 ± 0.02, p<2e-16, Table S20) after learning.

*Delta coherence*

When analyzing delta frequency coherence (1.5-4 Hz) we obtained comparable results to those obtained for SW coherence. The final stepwise reduced linear mixed model included a night effect (F(1, 5554.5) = 8.74, p= 0.003, Table S19,) and a night by genotype interaction (genotype x night: F(1, 5554.5) = 31.26, p= 2.36e-08). When estimating the marginal means using a least-squares approach we find increase in delta coherence in CC individuals (CC night 1: 0.78 ± 0.02, CC night 2: 0.79± 0.02, $p=0.003$, Table S20) but a reduction in those in the AA group (AA night 1: 0.81± 0.02, AA night 2: 0.78 ± 0.02, $p<2e-16$, Table S20).

*Slow sigma coherence*

Results for slow sigma coherence (9-12Hz) mimic those obtained for slow waves, where we see an overall night effect ($F(1, 5554.3) = 3.90$, $p= 0.048$, Table S19) and a genotype by night interaction ($F(1, 5554.3) = 12.42$, $p= 4.28e-4$). When estimating the marginal means using a least-squares approach we can detect an increase in slow sigma coherence in the CC group CC night 1: 0.72± 0.02, CC night 2: 0.73± 0.02 , $p= 0.0048$, Table S20) but a decrease in the AA group (AA night 1: 0.71± 0.02, AA night 2: 0.70 ± 0.02, $p = 0.002$, Table S20).

*Fast sigma coherence*

We found no evidence for main effects of genotype or night or interaction effects on fast sigma coherence (Table S19).

Overall, whole epoch NREM coherence effects are more pronounced compared to power and event amplitude effects, suggesting that synchronized NREM EEG rhythms support sleep dependent memory consolidation.

### Supplemental Discussion

#### NREM power

We observed consistent changes in NREM power in different frequency bands that were dependent on the combination of genotype and recording night. We observed consistent night effects, that underline the significance of learning dependent changes in cortico-thalamic oscillations that mark sleep-dependent memory consolidation. Although we observed predicted increases of slow and delta wave power in the CC group (non-risk) we found the opposite effect in the AA group (those carrying the SZ-associated allele), power in SW delta and slow spindles would decrease during the learning night.

Our results in the CC group are in line with studies showing increases in NREM oscillation density or amplitudes after learning in humans and animals. Previous studies showed an increase in SW/delta power (21), and increases in spindle density and amplitude (48–50) after motor learning in humans.

In our results changes in power seem mainly driven by changes in event amplitudes rather density of events (see main text for event-based analysis results, Tables S9-16) suggesting learning dependent changes in biophysical and network properties underlying slow/delta wave and spindle event amplitudes.

These signatures of sleep dependent memory processing seem impaired in carriers of the SZ-associated variant (AA group). To our knowledge no other studies have investigated learning dependent changes of sleep neurophysiology in carriers of common variants associated with schizophrenia, although a few recent studies described sleep phenotypes associated with genetic liability for SZ.

Schilling et al (51) demonstrated reduced that spindle activity is related to the dosage of (Val108) genotype in the gene catechol-O-methyltransferase (COMT). Homogenous carrier (Val/Val) show lower spindle density, which is in line with the fact that the Val/Val allele is conferring higher liability for SZ. In contrast, a recent study using polygenic risk scores for SZ in healthy participants, described a positive association between spindle amplitudes and density and polygenic risk score (52). i.e. higher risk was associated with higher spindle amplitudes. Our own results suggest that environmental factors may modulate differences in cortico-thalamic function during sleep in carriers of low liability/common variants.

Future studies using comparable experimental setups in larger cohorts should further delineate the interplay of genetic, environmental and plasticity mechanisms underlying sleep dependent memory processing in humans and its relation to psychiatric disorders.

#### NREM coherence

We also observed genotype and night dependent changes in frequency dependent connectivity during NREM sleep. Whole epoch NREM SW, delta and slow spindle coherence shows significant interaction effects. The CC group shows in increases in whole epoch SW, delta and slow spindle coherence, whereas the AA group shows a decrease in synchronization.

Previous studies have demonstrated increased coherence after learning during NREM sleep (53). Despite the fact, that overall global connectivity is reduced during NREM sleep in comparison to the wake state (54), brief epochs of synchronicity, orchestrated by SW and spindle oscillations may provide periods of increased neuronal functional co-activation during sleep that rely on long range anatomical connections (55–57).

Previous studies have shown decreased task dependent cortico-cortical coupling (58,59), and increased cortico-limbic functional connectivity (58,60,61) during wake in carriers of rs1344706. These observations are corroborated by deficits in grey and white matter structure in in carriers of the risk variant (62).

Our findings support and extend these early findings of task dependent functional connectivity in carriers of the SZ-associated variant to learning-dependent connectivity during NREM sleep. The deficit in coupling, in particular between frontal and occipital cortical areas during NREM may signify a deficit in coordinated re-activation during NREM sleep that is thought to underlie sleep dependent memory consolidation (63,64)

Although we did not identify obvious behavioral deficits, the increased variability in sleep dependent memory consolidation may point towards increased variability in corticothalamic development, network topologies and functional signatures. Since we recorded in a healthy population with a very small risk to develop a full psychiatric disorder it is likely that compensatory mechanisms protect against such small genetic liability. Future studies will aim to identify these compensatory mechanisms which may provide insights as how to achieve compensation of more severe sleep and cognitive phenotypes observed in patients diagnosed with psychiatric disorders.

### Supplemental Tables

#### Table S1 - Overview of statistical analysis of non-EEG data

| | Characteristic | Aim | Data source | Statistical approach | Results |
|---|---|---|---|---|---|
| (1) | Confounding factors | To compare possible confounding factors across genotype groups and between recruited and invited groups | Existing ALSPAC data | Categorical variables – Pearson Chi-square, Continuous variables - Wilcoxon rank-sum (Mann-Whitney) | Table S3 |
| (2a) | Motor Sequence Task performance | To compare overnight improvement in the MST across genotype groups | MST – overnight improvement in percent | Difference in means: Two-sample two-sided t-test (unequal variances);<br>Difference in variances: Two-sample variance comparison | Table 1, Table S21, Figure 2 |
| (2b) | Motor Sequence Task performance | To estimate the effect of genotype and session (training versus test) on MST performance | Average number of correct sequences and reaction times for last 3 trials (training, evening) and first 3 trials (test, morning). | Difference in means: Linear mixed model with MST performance as the dependent variable, genotype group and session as fixed effects and individual as a random effect,<br>Difference in variances: Levene's robust test for equality of variance | Table 2, Figure 2, Table S22 |
| (3a) | Diurnal sleep wake rhythms | To compare sleep behavior and daily rhythm across genotype groups | Actigraphy | Difference in means: Wilcoxon rank-sum (Mann-Whitney) | Table S4 |
| (3b) | Sleep architecture | To compare objectively measured sleep | PSG | Difference in means: Linear mixed model with sleep architecture variables as factors | Table S5 |
| (3c) | Subjective and objective sleep quality | To compare objectively measured sleep between those who self-reported good versus poor sleep quality | Pittsburgh Sleep Quality Index, actigraphy, PSG | Difference in means: Wilcoxon rank-sum (Mann-Whitney) | Table S6 |

**Table S2 - Overview of statistical analysis of sleep EEG data**

| | Characteristic | Aim | Data source | Statistical approach | Results |
|---|---|---|---|---|---|
| (1) | Spectral power of whole NREM epoch | To estimate the effect of genotype and night (first vs second) on spectral power and to explore possible interactions between the two factors | Noise free N2 & N3 epochs | Linear mixed model fitted using a stepwise reduction procedure, followed by predicted marginal means analysis. | Table S7, TableS8 |
| (2a) | Slow-wave event properties | To estimate the effect of genotype and night (first vs second) on event properties and to explore possible interactions between the two factors | NREM event properties derived from automatic detection during N2 & N3 | Linear mixed model fitted using a stepwise reduction procedure, followed by predicted marginal means analysis. | Table S9 Table S10, Figure 4 |
| (2b) | Delta wave event properties | | | | Table S11, Table S12 |
| (2c) | Slow spindle event properties | | | | Table S13 Table S14 |
| (2d) | Fast spindle event properties | | | | Table S15, Table S16, Figure 4 |
| (3a) | SW triggered slow coherence | To estimate the effect of genotype and night (first vs second) on SW triggered SW coherence and to explore possible interactions between the two factors | SW event triggered EEG data window (± 2s) for coherence analysis (N2 & N3) | Linear mixed model fitted using a stepwise reduction procedure, followed by predicted marginal means analysis. | Table S17, Table S18, Figure 5 |
| (3b) | Spectral coherence at SW frequency (0.5-1.5 Hz) of whole NREM epochs | To estimate the effect of genotype and night (first vs second) on NREM coherence and to explore possible interactions between the two factors | Noise free N2 & N3 epochs | Linear mixed model fitted using a stepwise reduction procedure, followed by predicted marginal means analysis. | Table S19, Table S20 |

**Table S3 - Characteristics of ALSPAC participants recruited and not recruited to the study**

| Potential confounders | | CC group (N=22) | | AA group (N=25) | | | Unrecruited (N=430) | | |
|---|---|---|---|---|---|---|---|---|---|
| | | N[a] | | N[a] | | *p*[b] | N[a] | | *p*[c] |
| Maternal education (% with A-level/Degree) | | 19 | 52.6% | 22 | 45.5% | 0.65[g] | 386 | 42.5% | 0.44[g] |
| Maternal social class (%) | | 16 | | 21 | | | 327 | | |
| | I - professional | | 18.8% | | 4.8% | 0.65[g] | | 8.9% | 0.29[g] |
| | II - managerial | | 31.3% | | 33.3% | | | 32.7% | |
| | III (non-manual) - skilled | | 37.5% | | 42.9% | | | 42.8% | |
| | III (manual) - skilled | | 12.5% | | 9.5% | | | 7.0% | |
| | IV – semi-skilled | | 0% | | 4.8% | | | 7.3% | |
| | V - unskilled | | 0% | | 4.8% | | | 1.2% | |
| PLIKS[d] questionnaire results at age 18 years (% with at least one symptom) | | 19 | 0 | 19 | 5.3% | 0.31[g] | 184 | 3.8% | 0.72[g] |
| WISC[e] Total IQ at age 8 (mean (SD)) | | 22 | 105.8 (18.7) | 23 | 107.0 (16.9) | 0.89[h] | 329 | 106.4 (15.7) | 0.97[h] |
| WASI[f] Total IQ at age 15 (mean (SD)) | | 19 | 95.2 (13.0) | 19 | 92.3 (10.6) | 0.31[h] | 234 | 93.2 (13.8) | 0.89[h] |

[a] N, no. of participants in the group with data for the outcome being tested; [b] test result from a comparison of CC to AA group; [c] test result from a comparison of recruited (N=47) to not recruited (N=430); [d] PLIKS, Psychosis-like symptoms; [e] WISC, Wechsler Intelligence Scale for Children; [f] WASI, Wechsler Abbreviated Scale of Intelligence; [g] Pearson Chi-square p-value; [h] Wilcoxon rank-sum (Mann-Whitney) test p-value.

**Table S4 - Habitual sleep behavior and diurnal rhythms derived from actigraphy**

| | Mean (SD) | | |
|---|---|---|---|
| | CC group | AA group | p[a] |
| Sleep analysis | N=16 | N=19 | |
| # nights in analysis | 14.3 (1.8) | 14.3 (1.7) | 0.75 |
| TIB (minutes) | 498 (31) | 490 (35) | 0.49 |
| TST (minutes) | 403 (35) | 393 (39) | 0.39 |
| SOL (minutes) | 10 (7) | 11 (8) | 0.86 |
| Sleep efficiency (%) | 81 (5) | 80 (6) | 0.62 |
| FI | 28.7 (8.4) | 27.8 (6.1) | 0.82 |
| NPCRA analysis | N=15 | N=19 | |
| # days in analysis | 13.3 (1.5) | 13.1 (1.9) | 0.79 |
| RA | 0.71 (0.08) | 0.73 (0.06) | 0.44 |
| IS | 0.68 (0.11) | 0.72 (0.12) | 0.21 |
| IV | 0.35 (0.07) | 0.34 (0.05) | 0.66 |
| L5 | 8.98 (2.51) | 8.12 (1.82) | 0.38 |
| M10 | 52.1 (4.0) | 52.5 (2.8) | 0.96 |

SD, standard deviation; TIB, time in bed; TST, total sleep time; SOL, sleep onset latency; FI, fragmentation index; RA, relative amplitude; IS, interdaily stability; IV, intra-daily variability; L5, least 5 average (L5); M10, most 10 average.

[a] *p*-value from Wilcoxon rank-sum (Mann-Whitney) test

#### Table S5 - PSG-derived sleep architecture across groups and sessions

| | Mean (SD) | | | | Mixed model output | |
|---|---|---|---|---|---|---|
| | CC group (N=18) | | AA group (N=22) | | Night effect | Genotype group effect |
| Outcome measure | Night 1 | Night 2 | Night 1 | Night 2 | Beta[a] (SE), F(df1,df2) , *p* | Beta[b] (SE), F(df1,df2) , *p* |
| **TIB (minutes)** | 498 (44) | 505 (47) | 497 (41) | 497 (40) | b = 2.89 (5.39), F = 0.29 (1, 39), p = 0.60 | b = -4.41 (12.42), F = 0.13 (1, 38), p = 0.72 |
| **TST (minutes)** | 457 (46) | 463 (47) | 448 (56) | 462 (43) | b = 9.91 (6.22), F = 2.54 (1, 39), p = 0.12 | b = -5.40 (14.11), F = 0.15 (1, 38) p = 0.70 |
| **Stage 1 %** | 7.3 (3.2) | 6.6 (3.0) | 8.2 (2.7) | 6.8 (2.3) | b = -1.09 (0.35), F = 9.47 (1, 39), p = $3.8 \times 10^{-03}$ | b = 0.50 (0.82), F = 0.37 (1, 38), p = 0.55 |
| **Stage 2 %** | 46.7 (7.1) | 42.2 (12.2) | 44.0 (7.7) | 44.2 (7.5) | b = -1.91 (1.27), F = 2.25 (1, 39), p = 0.14 | b = -0.37 (2.49), F = 0.02 (1, 38) p = 0.88 |
| **Stage 3 %** | 23.3 (6.3) | 23.5 (4.6) | 25.0 (6.1) | 25.4 (5.9) | b = 0.32 (0.51), F = 0.41 (1, 39)’ p = 0.52 | b = 1.81 (1.76), F = 1.05 (1, 38), p = 0.31 |
| **REM %** | 22.7 (3.8) | 24.7 (5.6) | 22.7 (5.7) | 23.7 (4.9) | b = 1.43 (0.71), F = 4.12 (1, 39), p = 0.05 | b = -0.54 (1.45), F = 0.14 (1, 38), p = 0.71 |
| **SOL (minutes)** | 17 (12) | 14 (9) | 12 (7) | 10 (5) | b = -2.67 (1.53), F = 3.06 (1, 39), p = 0.09 | b = -3.97 (2.20), F = 3.25 (1, 38), p = 0.08 |
| **WASO (minutes)** | 24 (12) | 28 (22) | 37 (26) | 25 (18) | b = -4.35 (4.34), F = 1.01 (1, 39), p = 0.32 | b = 4.96 (4.89), F = 1.03 (1, 38), p = 0.32 |
| **Sleep efficiency (%)** | 92 (4) | 92 (4) | 90 (6) | 93 (4) | b = 1.58 (0.91), F = 2.99 (1, 39), p = 0.09 | b = -0.35 (1.20), F = 0.09 (1, 38), p = 0.77 |

SD, standard deviation; TIB, time in bed; TST, total sleep time; SOL, sleep onset latency; WASO, wake after sleep onset.

[a] given with respect to night 1 as baseline; [b] given with respect to the CC group as baseline.

#### Table S6 - Sleep architecture in participants with PSQI scores above and below 5

| Mean (SD) | **PSQI <=5** | | **PSQI >5** | |
|---|---|---|---|---|
| | N=31 | | N=6 | |
| **PSG Architecture** | **Night 1** | **Night 2** | **Night 1** | **Night 2** |
| TIB (minutes) | 488 (33) | 493 (39) | 537 (62) | 534 (58) |
| TST (minutes) | 443 (43) | 459 (42) | 488 (77) | 484 (54) |
| SOL (minutes) | 14 (10) | 10 (6) | 12 (5) | 17** (3) |
| WASO (minutes) | 31 (20) | 25(18) | 36 (30) | 32 (16) |
| Sleep efficiency (%) | 91 (5) | 93 (4) | 91 (7) | 91 (3) |
| **Actigraphy** | **N=27** | | **N=6** | |
| TIB (minutes) | 488 (31) | | 513* (36) | |
| TST (minutes) | 396 (36) | | 412 (40) | |
| SOL (minutes) | 10 (6) | | 12 (11) | |
| Sleep efficiency (%) | 81 (5) | | 81 (6) | |

Asterisks indicate where there is evidence for a difference in means between the two groups (sleep architecture variable comparisons made within night): Wilcoxon rank-sum (Mann Whitney) test: * *p<0.05, **p<0.01.*

**Table S7 - Linear mixed model results for NREM power**

| | Stepwise reduced linear mixed model output | | | | | | | | |
|---|---|---|---|---|---|---|---|---|---|
| | **Night effect** | | | **Genotype effect** | | | **Interaction (Night X Genotype)** | | |
| | *Beta[c] (SE)* | *F(df1,df2)* | *p* | *Beta[c] (SE)* | *F(df1,df2)* | *p* | *Beta[c] (SE)* | *F(df1,df2)* | *p* |
| **Slow power (0.5-1.5 Hz)** | *0.22829 (0.08584)* | *7.0719 (1, 1373.03)* | *0.007921 ***** | *-0.45971 (0.52241)* | *0.7744 (1, 39.16)* | *0.384242* | *-0.38621 (0.12785)* | *9.1261 (1, 1373.03)* | *0.002566 ***** |
| **Delta power (1.5-4 Hz)** | *0.22160 (0.06171)* | *12.894 (1, 1373.02)* | *0.000343 ****** | *-0.19575 (0.46786)* | *0.175 (1, 38.74)* | *0.6779739* | *-0.38627 (0.09191)* | *17.663 (1,1373.02)* | *2.808e-05 ****** |
| **Slow sigma power (9-12 Hz)** | *0.21288 (0.08110)* | *6.8897 (1,1373.02)* | *0.0087656*** | *-0.14504 (0.69663)* | *0.0433 (1, 38.58)* | *0.8361661* | *-0.43461 (0.12078)* | *12.9477 (1,1373.01)* | *0.0003317 ****** |
| **Fast sigma power (12-15 Hz)** | *0.19811 (0.07725)* | *6.5763 (1,1373.02)* | *0.010440 ** | *0.05851 (0.69460)* | *0.0071 (1, 38.53)* | *0.933301* | *-0.30981 (0.11505)* | *7.2507 (1, 1373.01)* | *0.007174 ***** |

* p<0.05, ** p<0.01, ***p<0.001

#### Table S8 - Estimated marginal means of NREM power across all electrodes

| | Power (log((uV/Hz)$^2$) | | | |
|---|---|---|---|---|
| | **CC group (N=18)** | | **AA group (N=22)** | |
| | Night 1 | Night 2 | Night 1 | Night 2 |
| **Slow power (0.5-1.5 Hz)** | 19.8368 (0.3874) | 19.9947# (0.3874) | 20.6827 (0.3504) | 20.4544 (0.3505)** |
| **Delta power (1.5-4 Hz)** | 11.6892 (0.347) | 11.8538 (0.347)* | 12.2712 (0.3138) | 12.0496 (0.3139)** |
| **Slow sigma power (9-12 Hz)** | 0.3827 (0.5166) | 0.6044 (0.5166)* | 0.9623 (0.4673) | 0.7494 (0.4673)** |
| **Fast sigma power (12-15 Hz)** | -1.0223 (0.5151) | -0.9106 (0.5151) | -0.771 (0.466) | -0.9691* (0.466) |

Mean (SE), Within genotype, between night comparisons: # $p<0.1$, * $p<0.05$, ** $p<0.01$, *** $p<0.001$

#### Table S9 - Linear mixed model results for NREM SW event properties

| | Stepwise reduced linear mixed model output | | | | | | | | |
|---|---|---|---|---|---|---|---|---|---|
| | **Night effect** | | | **Genotype effect** | | | **Interaction (Night X Genotype)** | | |
| | Beta (SE) | F(df1,df2) | p | Beta (SE) | F(df1,df2) | p | Beta (SE) | F(df1,df2) | p |
| **SW density (N/min)** | 1.920e-02 (1.864e-02) | 1.0610 (1, 1373.18) | 0.3032 | 2.741e-02 (3.933e-02) | 0.4858 (1, 47.14) | 0.4892 | 8.772e-03 (2.515e-02) | 0.1216 (1, 1373.21) | 0.7273 |
| **SW amplitude (µV)** | 4.5681 (0.9865) | 21.4405 (1, 1356.04) | 4.00e-06 *** | 11.7354 (6.4694) | 0.8591 (1,38.81) | 0.0774 # | -5.7392 (1.3284) | 18.6669 (1, 1356.05) | 1.67e-05 *** |
| **SW frequency (Hz)** | -1.633e-03 (2.807e-03) | 0.3387 (1, 1356.45) | 0.5607 | -9.669e-03 (6.739e-03) | 2.0586 (1, 44.77) | 0.1583 | -8.985e-05 (3.779e-03) | 0.0006 (1, 1356.41) | 0.9810 |
| **SW length (duration, s)** | 9.012e-04 (2.425e-03) | 0.1381 (1, 1356.25) | 0.7102 | 3.105e-03 (7.616e-03) | 0.1662 (1, 41.76) | 0.6856 | -4.854e-04 (3.265e-03) | 0.0221 (1, 1356.23) | 0.8819 |

[grey box] = full model output, no evidence of night or genotype effect

**Table S10 - Estimated marginal means of NREM SW event properties across all electrodes:**

| | CC group (N=18) | | AA group (N=22) | |
|---|---|---|---|---|
| | Night 1 | Night 2 | Night 1 | Night 2 |
| **SW density (N/min)** | 0.6117 (0.0292) | 0.6309 (0.0291) | 0.6391 (0.0264) | 0.6671 (0.0264) |
| **SW amplitude (µV)** | 100.5537 (4.7984) | 105.1218 (4.7975)*** | 112.2891 (4.3392) | 111.118 (4.34) |
| **SW frequency (Hz)** | 1.1707 (0.005) | 1.1691 (0.005) | 1.1611 (0.0045) | 1.1593 (0.0045) |
| **SW length (duration, s)** | 0.7821 (0.0057) | 0.783 (0.0056) | 0.7852 (0.0051) | 0.7857 (0.0051) |

Mean (SE), within genotype, between night comparisons: # p<0.1, * p<0.05, ** p<0.01, ***p<0.001

**Table S11 - Linear mixed model results for NREM delta wave event properties**

| | Stepwise reduced linear mixed model output | | | | | | | | |
|---|---|---|---|---|---|---|---|---|---|
| | **Night effect** | | | **Genotype effect** | | | **Interaction (Night X Genotype)** | | |
| | Beta (SE), | F(df1,df2), | p | Beta (SE), | F(df1,df2), | p | Beta (SE), | F(df1,df2), | p |
| **Delta density (N/min)** | 5.019e-02 (3.007e-02) | 2.7872 (1, 1373.12) | 0.09525 | 4.796e-02 (7.697e-02) | 0.3883 (1, 43.88) | 0.53645 | -5.819e-02 (4.057e-02) | 2.0571 (1, 1373.14) | 0.15172 |
| **Delta amplitude (µV)** | 2.9055 (0.8777) | 10.9584 (1 1364.03) | 0.000956 *** | 8.7948 (5.4417) | 2.9055 (1 38.92) | 0.1141 | -3.6481 (1.1837) | 9.4984 (1, 1364.03) | 0.0020977 ** |
| **Delta frequency (Hz)** | 0.009 (0.0035) | 6.5674 (1,1365) | 0.01049 * | n.s. | n.s. | n.s. | n.s. | n.s. | n.s. |
| **Delta length (duration, s)** | 2.852e-03 (2.269e-03) | 1.5800 (1, 1364.13) | 0.2090 | 3.285e-03 (7.126e-03) | 0.2126 (1, 41.77) | 0.6472 | -1.403e-03 (3.060e-03) | 0.2103 (1, 1364.13) | 0.6466 |

[grey] = full model output, no evidence of night or genotype effect

**Table S12 - Estimated marginal means of NREM delta wave event properties across all electrodes**

| | **CC group (N=18)** | | **AA group (N=22)** | |
|---|---|---|---|---|
| | Night 1 | Night 2 | Night 1 | Night 2 |
| **Delta density (N/min)** | 1.1007 (0.0571) | 1.1509 (0.0571) | 1.1487 (0.0516) | 1.1407 (0.0516) |
| **Delta amplitude (µV)** | 92.1743 (4.0357) | 95.0798 (4.0355)*** | 100.9691 (3.6504) | 100.2266 (3.6504) |
| **Delta frequency (Hz)** | 2.2061 (0.0114) | 2.1978 (0.0114) | 2.1942$^{+}$ (0.0103) | 2.1846$^{+}$ (0.0103) |
| **Delta length (duration, s)** | 0.5571 (0.0053) | 0.5599 (0.0053) | 0.5604 (0.0048) | 0.5618 (0.0048) |

Mean (SE), within genotype, between night comparisons: # p<0.1, * p<0.05, ** p<0.01, ***p<0.001

**Table S13 - Linear mixed model results for NREM slow spindle event properties**

| | Stepwise reduced linear mixed model output | | | | | | | | |
|---|---|---|---|---|---|---|---|---|---|
| | **Night effect** | | | **Genotype effect** | | | **Interaction (Night X Genotype)** | | |
| | Beta (SE) | F(df1,df2) | p | Beta (SE) | F(df1,df2) | p | Beta (SE) | F(df1,df2) | p |
| **Slow spindle density (N/min)** | 0.03277 (0.03489) | 0.8824 (1, 1357.31) | 0.56 | 0.06791 (0.11473) | 0.3503 (1, 41.42) | 0.3477104 | -0.07445 (0.04712) | 2.4957 (1, 1357.36) | 0.1143902 |
| **Slow spindle amplitude (µV)** | 0.6354 (0.3302) | 4.8377 (1, 1331.06) | 0.054524 # | 2.7344 (2.7174) | 1.0125 (1, 38.52) | 0.320585 | -1.2918 (0.4451) | 8.4238 (1, 1331.07) | 0.003764 ** |
| **Slow spindle frequency (Hz)** | -4.674e-02 (1.646e-02) | 9.2693 (1,1332.3) | 0.0024 ** | n.s. | n.s. | n.s. | n.s. | n.s. | n.s. |
| **Slow spindle length (duration, s)** | 0.010 (0.0065) | 3.3330 (1,1331.81) | 0.068 | 1.591e-03 (2.251e-02) | 0.0050 (1, 45.30) | 0.94396 | 1.262e-02 (1.300e-02) | 0.9418 (1 1331.93) | 0.33200 |

= full model output, no evidence of night or genotype effect

**Table S14 - Estimated marginal means of NREM slow spindle event properties across all electrodes**

| | CC group (N=18) | | AA group (N=22) | |
|---|---|---|---|---|
| | Night 1 | Night 2 | Night 1 | Night 2 |
| **Slow spindle density (N/min)** | 0.5531 (0.0851) | 0.5858 (0.085) | 0.621 (0.077) | 0.5793 (0.077) |
| **Slow spindle amplitude (µV)** | 31.0691 (2.0154) | 31.7045 (2.0151) # | 33.8034 (1.8228) | 33.1471 (1.8229) * |
| **Slow spindle frequency (Hz)** | 11.3617 (0.0469) | 11.315 (0.0469) | 11.3283 (0.0424) | 11.3054 (0.0424) |
| **Slow spindle length (duration, s)** | 0.8749 (0.0167) | 0.8573 (0.0167) | 0.8733 (0.0151) | 0.8684 (0.0151) |

Mean (SE), within genotype, between night comparisons: # p<0.1, * p<0.05, ** p<0.01, ***p<0.001

**Table S15 - Linear mixed model results for NREM fast spindle event properties**

| | Stepwise reduced linear mixed model output | | | | | | | | |
|---|---|---|---|---|---|---|---|---|---|
| | **Night effect** | | | **Genotype effect** | | | **Interaction (Night X Genotype)** | | |
| | Beta (SE) | F(df1,df2) | p | Beta (SE) | F(df1,df2) | p | Beta (SE) | F(df1,df2) | p |
| **Fast spindle density (N/min)** | -4.142e-02 (4.174e-02) | 0.9844 (1, 1357.21) | 0.3213 | -1.264e-01 (1.660e-01) | 0.5797 (1, 40.29) | 0.4508 | 1.102e-02 (5.639e-02) | 0.0382 (1, 1357.24) | 0.8450 |
| **Fast spindle amplitude (µV)** | 0.8243 (0.3061) | 7.2541 (1, 1356.04) | 0.0072 ** | 2.3894 (2.6588) | 0.8076 (1, 38.46) | 0.3744 | -1.3232 (0.4136) | 10.2360 (1, 1356.05) | 0.0014 ** |
| **Fast spindle frequency (Hz)** | 3.897e-03 (1.145e-02) | 0.1157 (1, 1356.07) | 0.7338 | -9.165e-03 (8.061e-02) | 0.0129 (1, 38.71) | 0.9101 | -2.051e-02 (1.548e-02) | 1.7565 (1, 1356.08) | 0.1853 |
| **Fast spindle length (duration, s)** | 8.353e-03 (3.750e-03) | 4.9621 (1, 1356.42) | 0.026073 * | 2.466e-02 (1.060e-02) | 1.0824 (1, 42.73) | 0.02607 * | -1.363e-02 (5.067e-03) | 7.2399 (1, 1356.53) | 0.0072 ** |

= full model output, no evidence of night or genotype effect

#### Table S16 - Estimated marginal means of NREM fast spindle event properties across all electrodes

| | CC group (N=18) | | AA group (N=22) | |
|---|---|---|---|---|
| | Night 1 | Night 2 | Night 1 | Night 2 |
| **Fast spindle density (N/min)** | 1.7274 (0.1231) | 1.6859 (0.1231 | 1.601 (0.1114) | 1.5706 (0.1114) |
| **Fast spindle amplitude (µV)** | 31.8086 (1.9719) | 32.6329 (1.9717)* | 34.198 (1.7836) | 33.6991 (1.7837) |
| **Fast spindle frequency (Hz)** | 12.8534 (0.0598) | 12.8573 (0.0598) | 12.8442 (0.0541) | 12.8276 (0.0541) |
| **Fast spindle length (duration, s)** | 0.7946 (0.0079) | 0.8030 (0.0079)* | 0.8193 (0.0071)‡ | 0.8140 (0.0071) |

Mean (SE), within genotype, between night comparisons: # p<0.1, * p<0.05, ** p<0.01, ***p<0.001

Within night, between genotype effect: ‡ p<0.05

#### Table S17 - Linear mixed model results for SW triggered slow coherence

| | Mixed model output [b] | | | | | | | | |
|---|---|---|---|---|---|---|---|---|---|
| | **Night effect** | | | **Genotype effect** | | | **Interaction (Night X Genotype)** | | |
| | Beta[c] (SE) | F(df1,df2) | p | Beta[c] (SE), | F(df1,df2) | p | Beta[c] (SE) | F(df1,df2) | p |
| **SW triggered slow coherence (0.5-1.5 Hz)** | -3.305e-03 (2.838e-02) | 0.0455 (1, 11182) | 0.5128 | -3.305e-03 (2.838e-02) | 0.4365 (1, 38) | 0.8310 | 4.399e-02 (4.458e-03) | 97.3702 (1, 11182) | <2e-16 *** |

#### Table S18 - Estimated marginal means of SW triggered SW coherence across all electrodes

| | | | | |
|---|---|---|---|---|
| | **CC group (N=18)** | | **AA group (N=22)** | |
| | Night 1 | Night 2 | Night 1 | Night 2 |
| **SW triggered slow coherence (0.5-1.5 Hz)** | 0.8498916 (0.021) | 0.8714120 (0.019)*** | 0.8905784 (0.019) | 0.8681068 (0.021)*** |

mean(atanh(coherence)) (SE), within genotype, between night comparisons: # p<0.1, * p<0.05, ** p<0.01, ***p<0.001

**Table S19 - Linear mixed model results for whole epoch NREM coherence**

| | Stepwise reduced linear mixed model output | | | | | | | | |
|---|---|---|---|---|---|---|---|---|---|
| | **Night effect** | | | **Genotype effect** | | | **Interaction (Night X Genotype)** | | |
| | Beta[c] (SE), | F(df1,df2), | p | Beta[c] (SE), | F(df1,df2), | p | Beta[c] (SE), | F(df1,df2), | p |
| **Slow coherence (0.5-1.5 Hz)** | 1.131e-02 (5.025e-03) | 5.0623 (1, 5554.5) | 0.024 * | -0.005119 (0.029986) | 0.0291 (1, 39) | 0.8653 | -0.033651 (0.006788) | 24.5736 (1, 5554.4) | 7.365e-07 *** |
| **Delta coherence (1.5-4 Hz)** | 1.201e-02 (4.062e-03) | 8.7429 (1, 5554.3) | 0.0031 ** | 0.003568 (0.025354) | 0.0198 (1,38.9) | 0.8888 | -0.030676 (0.005486) | 31.2623 (1, 5554.4) | 2.362e-08 *** |
| **Slow sigma coherence (9-12 Hz)** | 9.583e-03 (4.848e-03) | 3.9070 (1, 5554.4) | 0.048 * | 0.026916 (0.033260) | 0.6549 (1, 38.7) | 0.4233078 | -0.023078 (0.006549) | 12.4186 (1, 5554.3) | 0.0004285 *** |
| **Fast sigma Coherence (12-15 Hz)** | -8.578e-04 (5.595e-03) | 0.0235 (1, 5554.3) | 0.8782 | -3.798e-02 (3.471e-02) | 1.1971 (1, 38.9) | 0.2806 | -5.670e-03 (7.557e-03) | 0.5629 (1, 5554.4) | 0.4531 |

(grey shading) = full model output with no evidence of night or genotype effect

* p<0.05, ** p<0.01, ***p<0.001

#### Table S20 - Estimated marginal means of whole epoch NREM coherence across all electrodes

| | atanh(coherence), mean (SE)[a] | | | |
|---|---|---|---|---|
| | **CC group (N=18)** | | **AA group (N=22)** | |
| | Night 1 | Night 2 | Night 1 | Night 2 |
| **Slow coherence (0.5-1.5 Hz)** | 0.848 (0.0222) | 0.8593 (0.0222)* | 0.8867 (0.0201) | 0.8644 (0.0201)*** |
| **Delta coherence (1.5-4 Hz)** | 0.7788 (0.0188) | 0.8059 (0.017)* | 0.7908 (0.0188) | 0.7872 (0.017)*** |
| **Slow sigma coherence (9-12 Hz)** | 0.7166 (0.0247) | 0.7262 (0.0247)* | 0.7128 (0.0223) | 0.6993 (0.0223)** |
| **Fast sigma Coherence (12-15 Hz)** | 0.8118 (0.0257) | 0.8109 (0.0257) | 0.7738 (0.0233) | 0.7673 (0.0233) |

Within genotype, between night comparisons: # p<0.1, * p<0.05, ** p<0.01, ***p<0.001

### Supplemental Figures

#### Figure S1 - CONSORT flow diagram of participant recruitment and exclusions prior to analysis

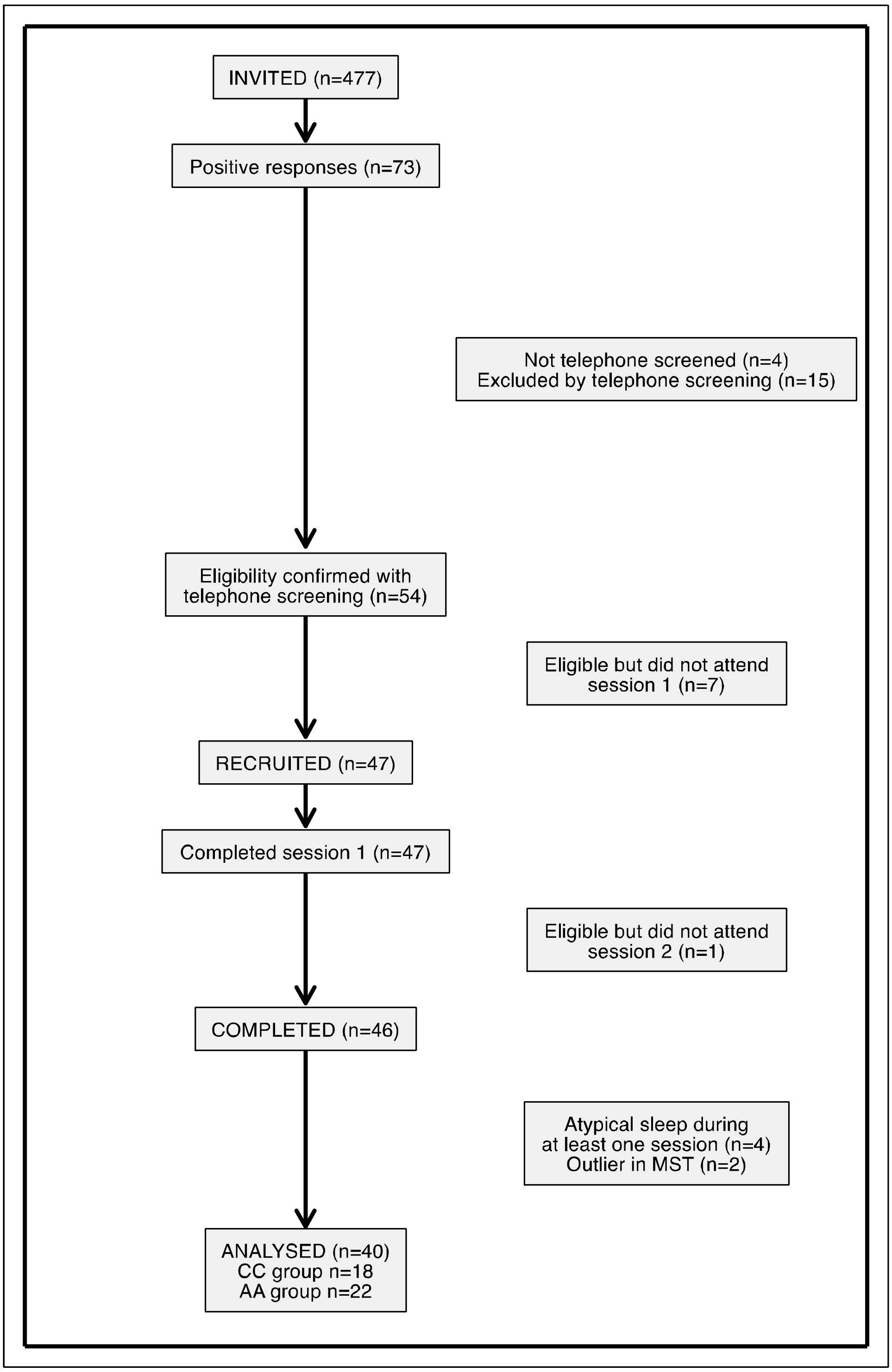
